# Herpes simplex virus infection promotes ALS pathology through ICP0-mediated PML body disruption

**DOI:** 10.64898/2026.03.27.714707

**Authors:** Dennis Freisem, Daniel Rombach, Sarah Brockmann, Anne Fink, Zoé Engels, Aurora de Luna, Dhiraj Acharya, Helene Hoenigsperger, Alicia Goreth, Sinje Tigges, Isabel Hagmann, Michiel van Gent, Fabian Zech, Anna Ponomarenko, Angela Rosenbohm, Johannes Dorst, Susanne Petri, Brit Mollenhauer, Jochen Weishaupt, Hayrettin Tumani, Mohamed R Gadalla, Daniela Huzly, Moritz Gaidt, Beate Sodeik, Abel Viejo-Borbolla, Markus Otto, Thomas Stamminger, Frank Kirchhoff, Adalbert Krawcyk, Ulf Dittmer, Lars Dölken, Tobias Böckers, Alberto Catanese, Gabriele Doblhammer, Georges MGM Verjans, Benedikt B Kaufer, Michaela U Gack, Florian Full, Hartmut Hengel, Veselin Grozdanov, Konstantin MJ Sparrer, Karin M Danzer

**Affiliations:** Institute of Molecular Virology, Ulm University Medical Center, Ulm, Germany; German Center for Neurodegenerative Diseases (DZNE), Ulm, Germany; Department of Neurology, Ulm University, Ulm, Germany; German Center for Neurodegenerative Diseases (DZNE), Bonn, Germany; Florida Research and Innovation Center, Cleveland Clinic, Port Saint Lucie, USA; Department of Viroscience, Erasmus Medical Center, Rotterdam, The Netherlands; Institute of Anatomy and Cell Biology, Ulm University, Ulm, Germany; Department of Child and Adolescent Psychiatry, Rostock University Medical Center, Rostock, Germany; Department of Neurology, Hannover Medical School, Hannover, Germany; Department of Neurology, University Medical Center Goettingen, Goettingen, Germany; Paracelsus-Elena Clinic, Centre of Parkinsonism and Movement Disorders, Kassel, Germany; Institute of Virology, Hannover Medical School, Hannover, Germany; Institute of Virology, University Medical Center and Faculty of Medicine, Albert-Ludwig-University Freiburg, Freiburg, Germany; National expert laboratory for HSV and VZV, University Medical Center, Freiburg, Germany; Research Institute of Molecular Pathology, Vienna BioCenter, Vienna, Austria; DZIF German Centre for Infection Research, Partner Site Hannover-Braunschweig, Hannover, Germany; Cluster of Excellence RESIST (EXC 2155), Hannover Medical School, Hannover, Germany; Department of Neurology, Martin-Luther-University Halle-Wittenberg, Halle (Saale), Germany; Institute of Virology, Ulm University Medical Center, Ulm, Germany; Institute for Virology, University Hospital Essen and University of Duisburg-Essen, Essen, Germany; Department of Infectious Diseases and Nephrology, University Hospital Essen, University Duisburg-Essen, Essen, Germany; Institute of Neuroanatomy, University Clinic RWTH Aachen, Aachen, Germany; Institute for Sociology and Demography, University of Rostock, Rostock, Germany; Institute of Virology, Freie Universität Berlin, Berlin, Germany

## Abstract

Transactive response DNA binding protein 43 kDa (TDP-43) pathology, is a central molecular hallmark of amyotrophic lateral sclerosis (ALS). However, the underlying triggers are incompletely understood. Here, we show that infection with herpes simplex virus (HSV) induces molecular hallmarks of ALS in various in vitro and in vivo models and is associated with an increased risk of ALS in human population data. German healthcare provider data (n = 238,440) and herpesvirus serology of an ALS patient and control cohort (n = 1,100) showed that HSV infection elevated the ALS risk by 210% and odds by ∼65%, respectively. On a molecular level, HSV infection promoted TDP-43 pathology in neuronal cell models, human iPSC-derived motoneurons and cerebral organoids, mice, and human tissue sections. This effect was triggered by HSV-1 or 2, but not by several other related herpesviruses. Mechanistically, the infected cell protein 0 (ICP0) of HSV-1/2 drives TDP-43 pathology by disturbance of promyelocytic leukemia nuclear bodies (PML-NBs), thereby abrogating TDP-43 SUMO2/3ylation. Taken together, we reveal a previously unrecognized association between HSV infection and ALS and clarify the underlying molecular mechanism that drives TDP-43 pathology. Our data may guide future studies into therapeutic and prophylactic interventions against ALS.

## INTRODUCTION

Amyotrophic lateral sclerosis (ALS) is a progressive neurodegenerative disorder (NDD) characterized by the selective degeneration of upper and lower motoneurons, leading to muscle weakness, paralysis, and ultimately death, typically within 2–5 years after symptom onset^1,2^. With 1 to 2.6 new cases per 100,000 people annually, ALS is a rare but devastating disease^2^. Only about 10-15% of ALS cases are linked to inherited Mendelian ALS mutations. Although familial ALS has revealed key pathological pathways, the etiology of sporadic ALS, which accounts for the majority of cases remains largely obscure, raising the possibility that environmental or exogenous factors may contribute to disease onset and progression. A unifying molecular hallmark of ALS pathology in more than 95% of all cases, is the mislocalization and aggregation of the RNA-binding protein TDP-43^3,4^. In healthy cells, TDP-43 is homogeneously distributed predominantly in the nucleus where it regulates RNA splicing, transcription, transport, and stability. Under physiological conditions TDP-43 continuously shuttles between the nucleus and cytoplasm, maintaining neuronal RNA homeostasis. Post-translational modifications regulate the solubility of TDP-43. For example, SUMO2/3ylation of TDP-43 in PML nuclear bodies (PML-NBs) promotes its solubility^5,6^. In contrast, pathological TDP-43 is frequently found as cytoplasmic, hyperphosphorylated, and highly ubiquitinated aggregates in affected neurons and glial cells^7^. These modifications stabilize misfolded protein species, promoting the formation of insoluble inclusions or pathological TDP-43 that can be processed to ∼35 kDa fragments, exacerbating toxic gain-of-function effects and further removal of TDP-43 from the nucleus. This functional depletion of TDP-43 from affected cells leads to widespread RNA-processing defects, including the aberrant expression of so-called cryptic exons that disrupt coding sequences across hundreds of mRNAs and neuronal death^8,9^. However, despite intensive research, the molecular mechanisms driving initial TDP-43 delocalization and aggregation remain poorly understood.

Emerging evidence suggests that viral infections contribute to the onset and progression of NDDs and other neuropathological diseases such as Alzheimer’s disease (AD), and multiple sclerosis (MS)^10–14^. One family of viruses that has been repeatedly associated with NDDs are herpesviruses, which include HSV-1 and 2, Varicella Zoster Virus (VZV) and Epstein-Barr Virus (EBV)^11,12,15^. These large DNA viruses establish lifelong latency in humans and are known to frequently infiltrate the central nervous system (CNS). For example, after initially replicating in the oral epithelium, HSV-1 enters sensory neurons and establishes latent infections in the trigeminal ganglia. From there it is able to reactivate to productive infections, leading to virus shedding with or without cold sores^16^. In rare cases infection of motoneurons may occur^17^. Virus dissemination upon reactivation and transmission into the CNS is thought to be less frequent, but associated with severe cases of encephalitis^18^. These reactivation events, often triggered by immunosuppression, UV light exposure or stress, may drive chronic neuroinflammation and neuronal injury^19^. EBV has been linked to MS in epidemiological studies, which showed that individuals infected with EBV are 32 times more likely to develop MS^20^. Mechanistical analyses suggest that the impact of EBV on MS may be mediated by auto-immune processes involving molecular mimicry and aberrant homing of T cells to the CNS^21^. Both HSV-1 and VZV reactivation have previously been linked to the development of dementias, including AD and vascular subtypes^13,22^. HSV-1 DNA and proteins have been detected in post-mortem human brains, particularly in regions vulnerable to AD pathology^23^. However, whether viruses play a role in the onset and progression of ALS remains unclear.

Here, we examined a possible association between herpesviruses and ALS by integrating epidemiological analyses, serological studies, *in vivo*– and molecular studies. Together, our data reveal a previously unrecognized link between HSV-1 and ALS. Infection with HSV-1 induces key molecular hallmarks of ALS in human induced pluripotent stem cell (iPSC)–derived cerebral organoids, and mouse models. Finally, our data shows that disruption of PML-NBs by the HSV-1 E3 ligase ICP0 drives the molecular pathology by preventing solubility-enhancing SUMOylation of TDP-43.

## RESULTS

### Population data show an association of HSV with ALS

To study the association of HSV infections and ALS, we analyzed retrospective incidence data in health-claim records from a national German healthcare provider. The nested-control, retrospective analysis comprising a 15-year follow-up used the records of 238,440 individuals aged ≥50 years without a definitive ALS diagnosis at the beginning of the observation period. During the follow-up period, 113 individuals were newly diagnosed with ALS (Criteria and full study design in Methods). The ALS patients were age and sex matched to 452 controls for a 1:4 ratio (Figure 1a; characteristics of the cohort in Supplementary Table 1). Until the ALS diagnosis/index date, 11.5% (n =13 of 113) of the case patients and 5.1% (n =23 of 452) of the controls had at least one HSV episode. The diagnosis was recorded as a B00 coding according to the ICD-10 medical coding system, which includes all forms of HSV-1/2 associated symptoms such as herpes blisters, nasal herpes, ocular herpes, herpes keratitis vesicular dermatitis, but also encephalitis or eczema. 7.1% of the ALS patients recorded HSV infections within 3 months (one calendar quarter) around the diagnosis, but only 3.5% of the controls. Infections were reported in 4.4% of the ALS patients and in 1.6% of the controls for two or more calendar quarters. Conditional logistic regression modelling adjusted for covariates (weighted Charlson comorbidity index, CCI) showed that having at least one HSV diagnosis is significantly associated with the risk of being diagnosed with ALS (odds ratio (OR)=3.1, 95% confidence interval (CI): 1.4-6.8, **p=0.005) (Fig. 1b). A one-unit increase in weighted CCI was associated with significantly increased odds of ALS (OR=1.1, 95% CI: 1.0-1.2, *p=0.046). Modelling the cumulative effect of multiple HSV diagnoses within prior quarters on the risk of ALS showed a 2.3-fold odds ratio increase for one HSV-1 diagnosis and ALS (OR=2.3, 95% CI: 0.9-5.8, p=0.07), but records of two or more HSV diagnoses were associated with a 7.6-fold odds ratio increase (OR=7.6, 95% CI: 1.5-538.1, **p=0.014) (Extended Data Fig. 1a). This suggests that multiple HSV diagnoses increase the likelihood of an ALS diagnosis. As a positive control, we used traumatic brain injury (TBI) and/or fractures of the skull and facial bones (ICD S06), which was previously associated with ALS^24^. In our data TBI increased the likelihood of ALS (OR=4.4, 95% CI: 1.0-18.9, p=0.07). Herpes zoster diagnosis, which was previously associated with dementia^13^, did not increase odds ratios for ALS.

**Fig. 1:**
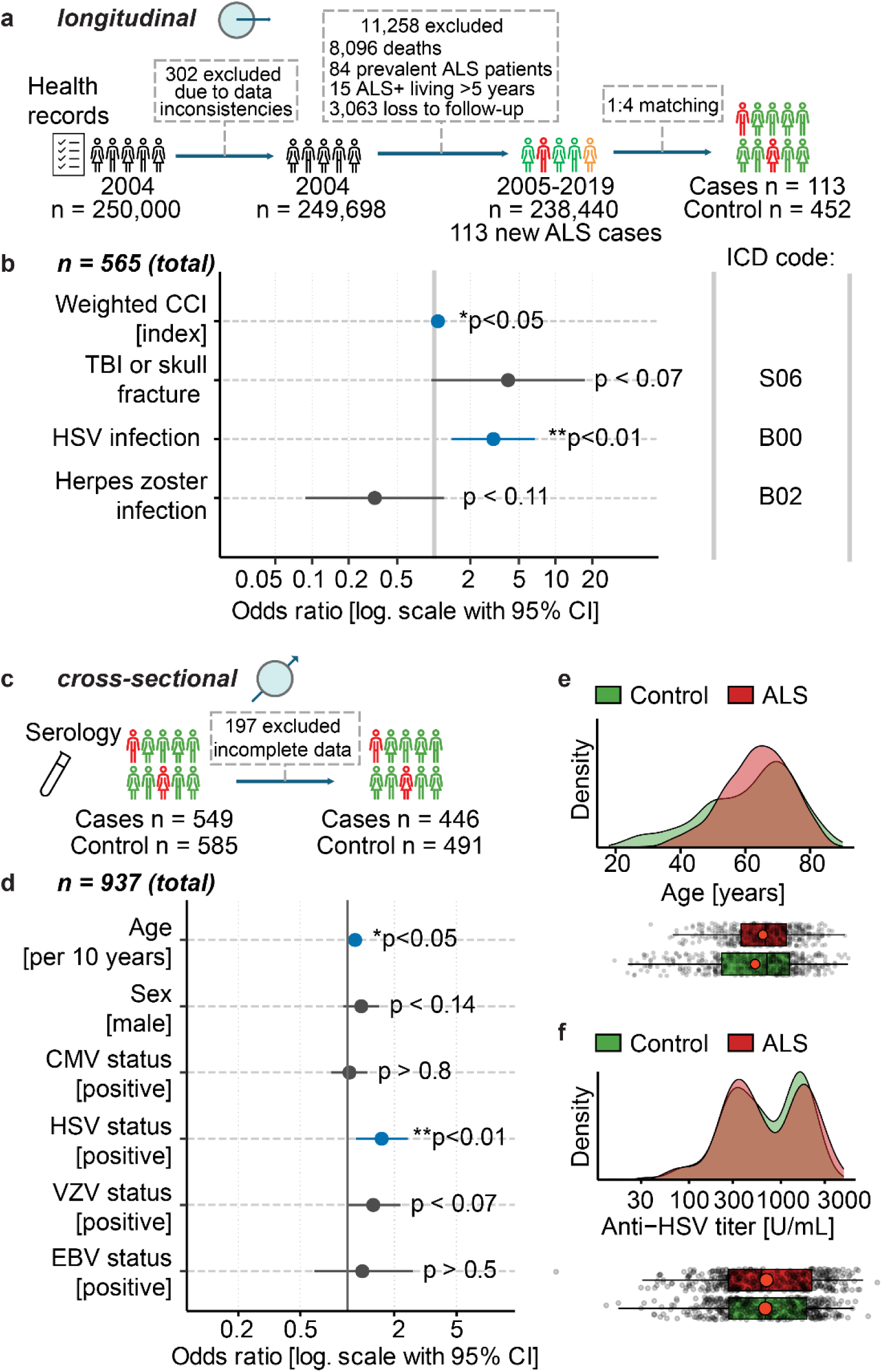
HSV-1 is associated with ALS. **a-b**, Epidemiological association of HSV-1 infections with ALS-diagnosis in a longitudinal, retrospective association study. The study design (a) included n = 250,000 individuals. n = 113 ALS diagnoses were recorded during the observation period and matched to n = 452 unaffected controls. ALS diagnosis was significantly associated with at least one prior HSV diagnosis (OR=3.1, 95% CI: 1.4-6.8, **p=0.005) (b). ICD: International Statistical Classification of Diseases and Related Health Problems, TBI: traumatic brain injury, CCI: Charlson Comorbidity Index. **c-f**, Epidemiological association of HSV-1 seropositivity with ALS diagnosis in a cross-sectional cohort of n =937 individuals. ALS diagnosis was significantly associated with seropositivity for HSV (OR: 1.65, 95% CI: 1.1-2.4, **p=0.009) and age (OR: 1.12, 95% CI: 1.0-1.3, *p=0.035 per 10 years of age) in a multiple logistic regression model (d). Age distribution of the ‘ALS’ and ‘Control’ groups (e) and HSV titer distribution in both groups (f). Box plots with median (center line), first and third quartiles (25/75%) and minimum/maximum value within 1.5x interquartile range from the first/third quantile, respective (lower/upper whisker). CMV: human cytomegalovirus, VZV: Varizella Zoster virus, EBV: Epstein-Barr virus.

These data suggest that HSV-1 diagnosis is associated with ALS diagnosis in population data.

### Positive serology of HSV-1 is associated with ALS

To corroborate the longitudinal analysis, we conducted a cross-sectional serological analysis of HSV prevalence in ALS patients and healthy controls. We assessed the antibody status against multiple herpesviruses (HSV-1/2, VZV, EBV, and cytomegalovirus (CMV)) in a cohort of 446 sporadic ALS patients, matched with 491 neurologically unaffected, age- and sex-matched controls from multiple centers in Germany (Fig. 1c, e). As expected, ∼60% (n = 611) of the individuals in the cohort tested positive for CMV antibodies, 85% (n = 953) positive for HSV-1/2 antibodies, ∼90% (n = 839) for VZV antibodies, and 96% (n = 1,089) for EBV antibodies (Extended Data Fig. 1b). Overall, for 83% of the individuals, the full antibody panel was completed, and they were included in the subsequent analyses (n = 937). As a control, the analyses showed that ALS was significantly associated with age which is a known risk factor for ALS (OR: 1.12, 95% CI: 1.0-1.3, *p=0.035 per 10 years of age, Fig. 1d). Interestingly, multiple logistic regression revealed that only the HSV serostatus was associated with a significant 1.6-fold increase in odds of ALS (OR: 1.65, 95% CI: 1.1-2.4, **p=0.009). Of note, similar titers of HSV-1/2 serum antibodies were observed in ALS patients and controls (Fig. 1f). None of the other tested herpesviruses was associated with a significant increase in risk of developing ALS. Finally, no correlation with past, non-herpesviral infections (influenza, measles, lassa and chikungunya virus) and ALS was observed (Extended Data Fig. 1c).

Taken together, the serology data support a significant association between HSV-1/2 and ALS.

### HSV-1 infection triggered delocalization of TDP-43

A molecular hallmark of ALS is the delocalization of TDP-43 from a uniform nuclear distribution into punctate-like structures, observed in the nucleus and cytoplasm^4^. Confocal microscopy analyses of overexpressed venus-YFP-tagged TDP-43 (TDP-43venus) in uninfected H4 neuroglioma cells showed that TDP-43 was almost uniformly distributed across the nucleus, which is consistent with published data (Fig. 2a)^4^. In contrast, infection with HSV-1 promoted a marked TDP-43 delocalization in the infected cells associated with a significant increase in overall numbers of TDP-43 puncta, indicating condensation of TDP-43 (Fig. 2a, b). Imaging analyses revealed that in uninfected H4 cells, TDP-43venus was localized mainly to the nucleus. However, upon infection with HSV-1 the TDP-43 signal in the cytoplasm increased significantly (Fig. 2c, d). Fractionation assays corroborated that both venus-tagged and endogenous TDP-43 were found predominantly in the nucleus of uninfected cells, but shifted towards a more cytoplasmic localization upon HSV-1 infection (Extended Data Fig. 2a, b). Inhibition of HSV-1 entry using neutralizing antibodies abrogated the puncta accumulation of TDP-43 as assessed by live cell imaging and automated puncta quantification (Extended Data Fig. 2c, d). To determine whether this delocalization was specific to TDP-43 and not a general effect on nuclear splicing or transcription factors, we infected H4 cells expressing a GFP-tagged version of the human serum response factor (hSRF) with HSV-1. Our analyses showed that TDP-43, but not hSRF, accumulated in puncta upon HSV-1 infection (Extended Data Fig. 2e). To probe whether puncta formation was specific and not due to stress induced by any infection, we infected cells with HSV-1, HSV-2, VZV, HCMV, or the non-herpesviral measles virus (MeV) and encephalomyocarditis virus (EMCV). Live-cell imaging aided automated image analyses showed that the number of TDP-43 puncta significantly increased over time during HSV-1/2 infection, but not following VZV, HCMV and MeV infection (Extended Data Fig. 2f). Upon EMCV infection, the strong cytopathic effect led to a rapid decrease of the TDP-43venus signal. Infection rates for the different viruses were similar in these experiments, as illustrated by immunofluorescence analyses (Extended Data Fig. 2g).

**Fig. 2:**
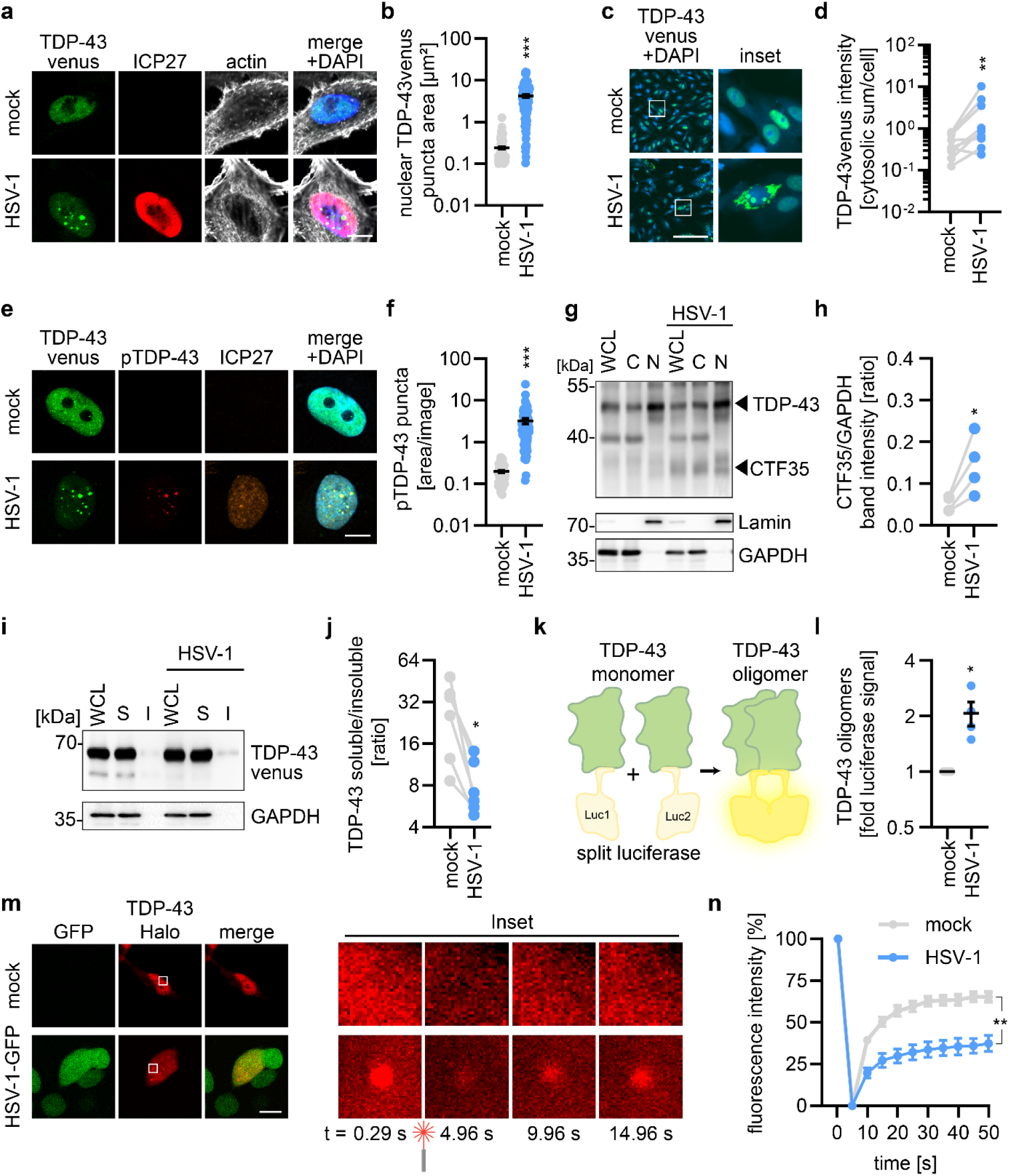
HSV-1 infection induces molecular hallmarks of ALS in cell culture models. **a**, Immunofluorescence of H4 cells overexpressing TDP-43venus (green) via lentivirus transduction were after 3 days, infected with HSV-1 (MOI 0.3). Cells were fixed after 17 h. ICP27 (red), DAPI (blue), actin (white). Scale bar, 10 µm. **b**, Analysis of nuclear TDP-43venus foci area in (**a**) using foci analyzer. Lines represent mean±SEM, n = 101-110, three independent biological replicates. ***p<0.001, t test. **c**, Representative immunofluorescence of H4 cells transduced with AAV expressing TDP-43venus (green). 3 days post-transduction, cells were infected with HSV-1 (MOI 0.3). 17 h post-HSV-1 infection, cells were fixed. DAPI (blue). White squares indicate origin of inset. Scale bar, 200 µm. **d**, Quantification of the total cytosolic TDP-43venus signal divided by the cell number per image as assessed by DAPI count, using Imaris software. Dots represent individual replicates. n = 9, three independent biological replicates. **p<0.01, paired t test. **e**, Representative immunofluorescent images of H4 cells transduced with lentiviruses expressing TDP-43venus (green) and 3 days later, infected with HSV-1 (MOI 0.3). Cells were fixed 17 h post-HSV-1 infection and stained for pTDP-43 (red), ICP27 (orange), and DAPI (blue). Scale bar, 10 µm. **f**, Quantification of the pTDP-43 area after thresholding for pTDP-43 dots (ImageJ). Lines represent mean±SEM, n = 60, three independent biological replicates. ***p<0.001, t test. **g**, Representative immunoblot of TDP-43 and GAPDH in H4 cells transduced with AAV expressing TDP-43venus for 3 days and infected with HSV-1 for 17 h. Cells were harvested and separated into whole cell lysate (WCL), nuclear (N) and cytosolic (C) fractions. Stained with anti-TDP-43, anti-Lamin, and anti-GAPDH. **h**, Quantification of the 35 kDa C-terminal fragment (CTF35) WCL band intensity in (**g**). Normalized to respective GAPDH band. n = 4, *p<0.05, paired t test. **i**, Representative immunoblots of HEK293T cells transduced with AAV-TDP-43venus (3 d) and infected with HSV-1 (MOI 0.3, 17 h). Fractions of the whole cell lysate (WCL) were separated into soluble (S) and insoluble (I) components. **j**, Western blot band intensities of soluble and insoluble TDP-43venus in (**i**) displayed as ratio. n = 6, *p<0.05, paired t test. **k**, Schematic representation of the TDP-43 split luciferase system. Upon TDP-43 oligomerization the fusion of L1 and L2 results in a functional luciferase. **l**, Quantification of luciferase activity in HEK293T cells co-transfected with split luciferase: TDP43-L1 and TDP43-L2. 24 h post-transfection cells were infected with HSV-1 (MOI 0.3). After 48 h post-infection cells were lysed and luciferase activity was analyzed. Bars represent mean±SEM of n = 4. *p<0.05, t test, **m**, Fluorescence recovery after photobleaching (FRAP) of H4 cells transduced with lentiviruses expressing TDP-43halo (red) and infected with HSV-1 GFP. 17 h post-infection, GFP positive cells with TDP-43halo delocalization were selected and analyzed. High laser intensity (100%) was applied to the region of interest (ROI) for 5 s. Subsequently, 10 images were acquired every 5 s. Scale bar, 20 µm. **n**, Quantification of the fluorescence recovery in ROIs as shown in (**m**). Pre-bleaching mean fluorescence intensity (MFI) was set to 100% and post-bleaching MFI was set to 0%. n = 12 individual cells. A linear mixed model with time and treatment as fixed effects and replicate (n = 9) as a random effect demonstrates a significant (**p=0.00272) effect for treatment and treatment:timepoint interaction (***p=2.45 x 10^-8^).

These results indicate that infection specifically with HSV-1 or HSV-2, but not with other viruses, promote an ALS-typical condensation of TDP-43 into puncta and delocalization into the cytoplasm.

### HSV-1 promotes processing, phosphorylation, oligomerization, and phase separation of TDP-43

Other hallmarks of ALS pathology include the (hyper)phosphorylation, proteolytic processing, decreased solubility, oligomerization, and phase-separation of TDP-43 into aggregate-like puncta^4,25^. Immunofluorescence images showed elevated levels of phosphorylated TDP-43 in HSV-1–infected H4 cells (Fig. 2e, f). False-positive signals from pTDP-43 antibody binding by the viral Fc receptor gI/gE of HSV-1 were excluded (Extended Data Fig. 2h). ALS-associated processing of TDP-43 from its full-length (43 kDa for endogenous TDP-43) to a smaller (35 kDa) C-terminal fragment, was monitored by western blotting (Fig. 2g). HSV-1 infection of H4 cells significantly increased the relative amount of the processed C-terminal fragment of endogenous TDP-43 (Fig. 2h). To assess whether HSV-1 affected TDP-43 solubility, we performed (in)solubility assays by biochemically fractioning. These results showed a markedly reduced ratio of soluble to insoluble TDP-43venus upon HSV-1 infection, consistent with enhanced protein aggregation (Fig. 2i, j). This was further supported by an oligomerization assay based on protein-complementation^26^. TDP-43 was fused to split Gaussia luciferase fragments, thus TDP-43 in close proximity complements the reporter leading to bioluminescence activity as a proxy for oligomerization (Fig. 2k). HSV-1 infection led to a robust increase in TDP-43 oligomerization (Fig. 2l). Finally, to probe the phase separation of TDP-43 puncta, we performed fluorescence recovery after photobleaching (FRAP) experiments using Halo-tagged TDP-43 in H4 cells. In uninfected H4 cells, the TDP-43-Halo signal recovered rapidly in the bleached areas (Fig. 2m, n). In contrast, the HSV-1-induced TDP-43 puncta exhibited a significantly slower recovery. In addition, while non-punctate TDP-43 recovered ∼75% of the pre-photobleaching signal intensity within 50 s, the infection-induced puncta reached only below 50% of its original intensity. This indicates the formation of intracellular TDP-43 assemblies with reduced mobility.

Taken together, these assays demonstrate that HSV-1 promotes key molecular hallmarks of ALS in infected cells, including TDP-43 delocalization, processing, oligomerization and phase separation.

### HSV-1 infection delocalizes TDP-43 in physiologically relevant *in vitro* and *in vivo* models

To assess HSV-1 induced TDP-43 pathology in more relevant systems we used 2D *in vitro* cultures (primary mouse cortical neurons and human induced pluripotent stem cell (iPSC)-derived motoneurons), complex 3D cultures (iPSC-derived human forebrain organoids) and *in vivo* data (mice and human tissue sections). Primary cortical neurons were transduced with adeno-associated virus (AAV) expressing TDP-43venus and infected with HSV-1. The infection caused a puncta-like appearance of TDP-43 and shifted its localization towards the cytoplasm (Fig. 3a, b; Extended Data Fig. 3a). To analyze the impact of HSV-1 infection in a 2D human neuron model, we differentiated motoneurons from human iPSCs and infected the cultures with HSV-1. While endogenous TDP-43 was distributed in the nucleus of uninfected cells, in the infected cells TDP-43 puncta appeared (Fig. 3c, d).

**Fig. 3:**
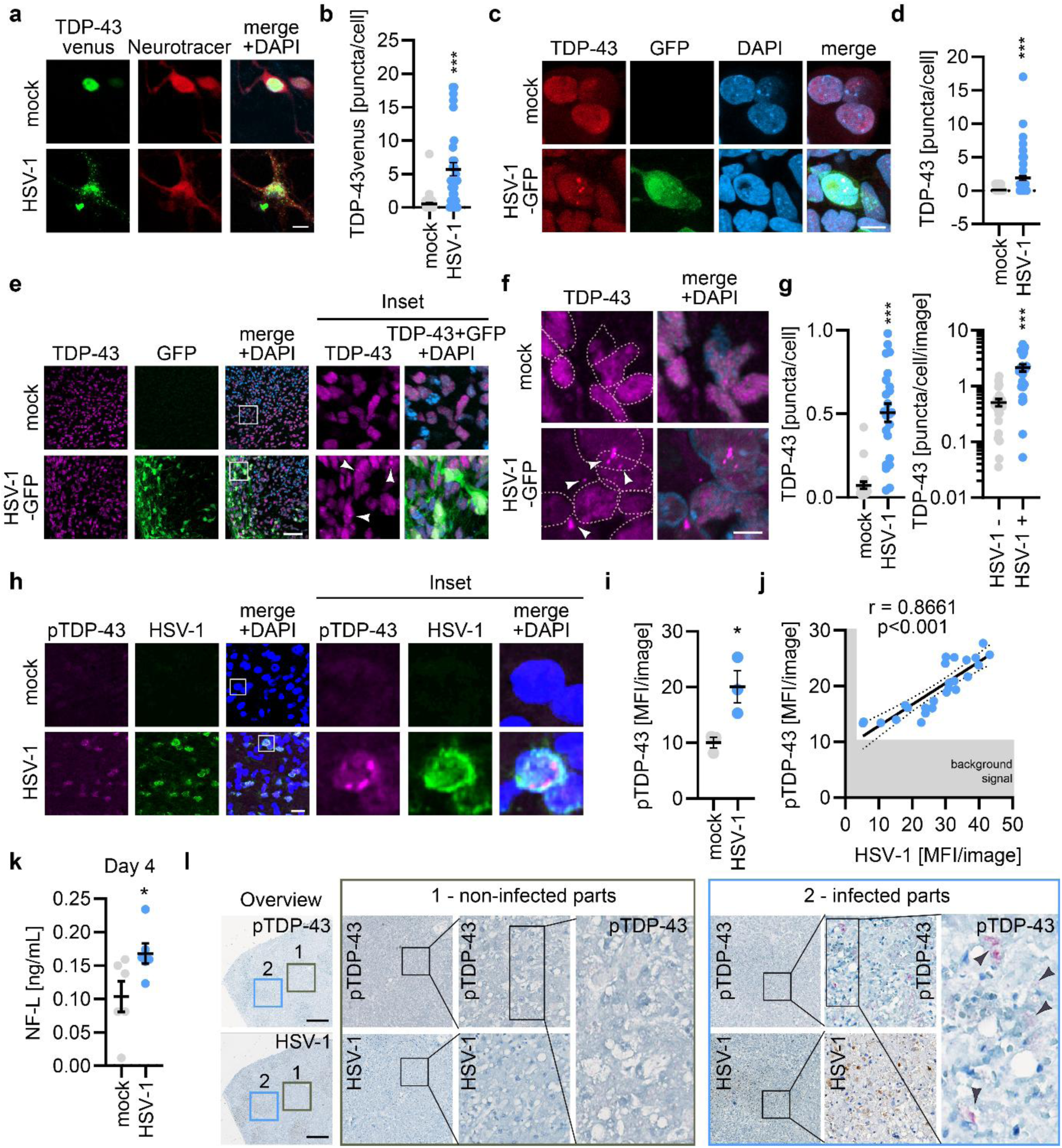
HSV-1 infection delocalizes TDP-43 in complex models and *in vivo*. **a**, Primary murine neurons were isolated from C57BL/6J mice at postnatal day 0–1. Neurons were transduced with AAV expressing TDP-43venus (green). After 3 days, cells were infected with HSV-1 and 17 h later fixed. Neurotracer (red), DAPI (blue). Scale bar, 10 µm. **b**, Analysis of TDP-43venus puncta per cell in (**a**). n = 31-43, three biological replicates. ***p<0.001, t test. **c,** Three-week-old human iPSC derived motoneurons (MNs) were HSV-1 GFP (green) infected and 17 h later stained for TDP-43 (red). DAPI (blue). Scale bar, 10 µm. **d**, Quantification of TDP-43 puncta in HSV-1 or mock infected MNs in (**c**). n = 64-68, two biological replicates. ***p<0.001, t test. **e**, Human iPSC derived brain organoids infected with HSV-1 GFP (green) for 48 h. Organoid slices were stained for TDP-43 (magenta), DAPI (blue). White arrows indicate TDP-43 delocalization in HSV-1 GFP positive cells. White squares indicate origin of inset. Scale bar, 50 µm. **f**, Zoom-in of representative cells in either condition in (**e**), white arrow indicate perinuclear TDP-43. Nuclei are framed by the dotted line. Scale bar, 5 µm. **g**, Quantification of TDP-43 puncta per cell in infected (HSV-1) and uninfected (mock) organoids (left) in (**e**, **f**), and analysis of TDP-43 puncta per cell per image in HSV-1 GFP infected organoids in GFP negative (HSV-1 −) or GFP positive (HSV-1 +) cells (right). n = 21-24, two independent replicates of two donors, ***p<0.001, t test. **h**, Representative immunofluorescence staining of C57BL/6J mice (10-week-old) that were intracerebrally infected with HSV-1 (1 x 10^5^ PFU/mice) or inoculated with PBS (mock). At day 3 post-infection, mid-sagittal sections were prepared and stained for phosphorylated TDP-43 (pTDP-43, magenta), HSV-1 (green), and nuclei (DAPI, blue). White squares indicate origin of inset. Scale bar, 20 µm. **i**, Quantification of pTDP-43 mean fluorescence intensity (MFI) in 8-10 unbiasedly selected brain areas for each mouse. n = 3 mice per group, *p<0.05, t test. **j**, Scatter plot of pTDP-43 MFI/image (y-axis) against HSV-1 MFI/image (x-axis). Each dot represents one corresponding measurement from HSV-1 infected mice in (**i**). Grey areas represent averaged MFI in mock infected mice (background signal). The line indicates the linear regression including 95% confidence intervals. Spearman’s rank correlation test (r = 0.8661, ***p<0.001, n = 25). **k**, NF-L serum levels of C57BL/6J mice (16-week-old) at day 4 post-intracerebral HSV-1 infection (5 x 10^5^ PFU/mice) or mock treatment, determined by ELISA. n = 6 mice per group, *p<0.05, t test. **l**, Representative immunohistochemistry of human liver section from an HSV-1 hepatitis patient. HSV-1 stained in brown, pTDP43 stained in magenta. Scale bar, 500 µm. Insets are indicated of infected (HSV-1 positive cells, brown) and non-infected areas. Arrows indicate pTDP-43 accumulations.

Next, to mimic the 3D architecture as well as to provide a more complex cell type composition we used iPSC-derived forebrain organoids. Organoids were differentiated for 90 days and contain cortical neurons, astrocytes and oligodendrocytes^27^. As expected, endogenous TDP-43 was mainly localized in the nucleus of the cells (Fig. 3e, f). However, upon infection of the organoids with an HSV-1 reporter strain expressing GFP, TDP-43 was redistributed to puncta-like structures in the nuclear and peri-nuclear regions of the infected cells (Fig. 3e, f). Quantification confirmed a significant increase of TDP-43 puncta in the nuclei of infected brain organoid sections (Fig. 3g). Of note, within a slice, TDP-43 delocalization was largely restricted to the HSV-1 infected cells (Fig. 3g). False-positive reaction of the used antibodies targeting endogenous TDP-43 with the viral Fc receptor was excluded (Extended Data Fig. 3b, c).

To determine whether HSV-1 infection also causes TDP-43 pathology *in vivo*, we investigated the phosphorylation status of TDP-43 in HSV-1 infected mice. C57BL/6J mice were either mock-inoculated or intracerebrally infected with HSV-1. At day 3 post-infection, the mice were sacrificed and their brain sections analyzed by immunofluorescence. False positive reaction of the used antibodies with the viral Fc receptor was excluded (Extended Data Fig. 3d). As opposed to the brains of mock-inoculated mice, the brain of HSV-1 infected mice showed a clear punctate pTDP-43 signal that was significantly more intense (Fig. 3h, i). Of note, the pTDP-43 mean fluorescence intensity correlated with the HSV-1 signal (Pearson’s correlation, r = 0.87, p <0.0001), indicating that more robust viral replication leads to higher pTDP-43 levels (Fig. 3j). Concurrently, serum levels of neurofilament light chain (NF-L), a biomarker of ALS^28^, in mock-treated and HSV-1 infected mice were analyzed by ELISA. The NF-L levels were significantly elevated at day 4 post-infection in the HSV-1 infected group (as compared to mock-treated mice) (Fig. 3k). Notably, at this time the infected mice exhibited only minor weight loss and did not show overt disease (Extended Data Fig. 3e). To corroborate these findings in human tissue, we analyzed liver tissue sections from a patient with a rare HSV-1 liver infection and compared them to a patient who was infected with hepatitis B virus (HBV). Notably, immunohistochemistry analyses using diagnostic pTDP-43 staining revealed that areas positive for HSV-1 show increased pTDP-43 staining as aggregate or skein-like formations in infected areas (Fig. 3l). In contrast, liver tissue from a patient with an HBV infection, which was stained in parallel, showed no discernable pTDP-43 signal (Extended Data Fig. 3f).

Taken together, our *in vitro* and *in vivo* studies confirm that HSV-1 infection promotes ALS-associated delocalization and phosphorylation of TDP-43 in primary mouse neurons, human motor neurons, human forebrain organoids, mouse models and human tissue slices.

### HSV-1 ICP0 gene expression drives TDP-43 delocalization

Gene expression of HSV-1 is regulated in a temporal order of three transcriptional continuums, immediate early (IE), early (E) and late (L) (Fig. 4a)^29^. The five IE genes (ICP0, ICP4, ICP22, ICP27, ICP47) are expressed already within the first few hours after infection, followed by ∼25 early genes, and subsequently late expression of >45 genes commences after viral DNA replication begins. To understand which gene expression phase triggers TDP-43 delocalization, we performed live-cell imaging analysis in H4 cells. After HSV-1 infection, TDP-43venus puncta were detected as early as ∼4 h post-infection (Fig. 4b, c). Similarly, the average object size of the TDP-43venus signal decreased ∼4 h post-infection suggesting a condensation into smaller structures (Extended Data Fig. 4a, b). Tomography live-cell imaging microscopy revealed that the transition of TDP-43venus from nuclear distribution to intranuclear puncta occurs rapidly within 15-30 min of HSV-1 infection (Fig. 4d). Acyclovir effectively blocks viral DNA replication and late gene expression^30^. TDP-43venus delocalization and puncta formation were still apparent 3 days post-infection and upon treatment with acyclovir (7h post-infection) (Fig. 4e). Time-resolved analyses further showed that even treatment with acyclovir as early as 4 h post-infection did not effectively prevent TDP-43 delocalization (Fig. 4f). This suggested that late gene expression does not contribute to TDP-43 delocalization. Combined with the rapid onset of TDP-43 particle formation, following HSV-1 infection, we hypothesized that HSV-1 gene products from the immediate early phase —ICP0, ICP4, ICP22, ICP27, and/or ICP47—mediate these alterations. To identify the responsible viral protein, H4 cells were co-transfected with TDP-43venus and plasmids expressing individual IE genes. Confocal imaging and quantitative analysis of nuclear TDP-43venus foci area revealed that expression of HSV-1 ICP0 alone, but not of the other HSV-1 proteins, was sufficient to induce robust TDP-43venus delocalization to a similar extent as HSV-1 infection (Fig. 4g, h). Overall, the mean fluorescence intensity of TDP-43venus was not reduced by co-expression of any tested viral protein, while ICP0 expression even slightly increased the signal (Extended Data Fig. 4c). To determine whether ICP0 was also required in the viral context to induce TDP-43 puncta formation, we used recombinant HSV-1 that lacked expression of ICP0 (HSV-1ΔICP0) and infected H4 cells expressing TDP-43venus. Live-cell imaging analyses coupled to automated TDP-43venus puncta counting revealed significantly less TDP-43venus puncta formation upon infection with HSV-1ΔICP0 (Fig. 4i, j, Extended Data Fig. 4d) despite similar infection levels (Extended Data Fig. 4e).

**Fig. 4:**
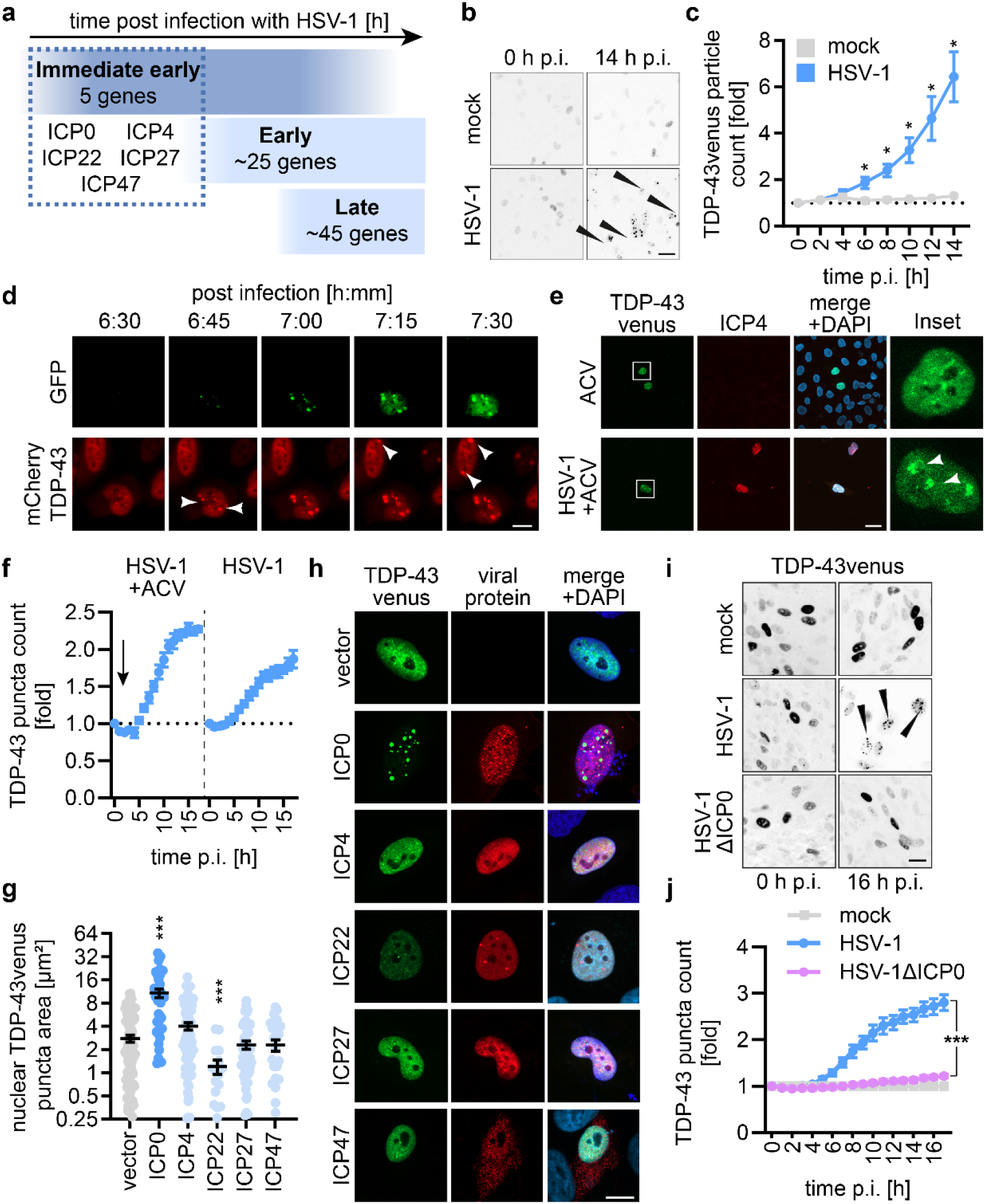
HSV Immediate early protein ICP0 drives TDP-43 pathology. **a**, Schematic illustration of HSV-1 protein expression after infection over time, with 5 immediate early genes (ICP0, ICP4, ICP22, ICP27 and ICP47), around 25 early and 45 late genes. **b**, Representative endpoint fluorescence images of H4 cells transduced with AAV overexpressing TDP-43venus and after 3 days, infected with HSV-1 (MOI 0.3). Every 2 h an image was acquired for 14 h post-HSV-1 infection. Black arrows indicate delocalized TDP-43venus. Scale bar, 20 µm. **c**, Quantification of TDP-43venus particles in (**b**), normalized on particle count at 0 h post-infection for respective condition. Dotted black line represents the base line (y = 1). Dots represent the mean±SEM of n = 3, three independent biological replicates. *p<0.05, t-test. **d**, Representative fluorescence images of H4 cells transduced with lentiviruses expressing TDP-43mCherry (red) and infected 3 days post-transduction with HSV-1 GFP (green) (MOI 0.3). Images were acquired every 15 min post-HSV-1 GFP infection using the 3D Cell Explorer-fluo (NanoLive). White arrows indicate TDP-43mCherry delocalization. Scale bar, 10 µm. **e**, H4 cells transduced with lentivirus expressing TDP-43venus (green) for 3 days were infected with HSV-1 (MOI 0.3). Acyclovir (ACV) was added to the cells 7 h post-infection (1 mM). Cells were fixed 3 days post-infection. ICP4 (red), DAPI (blue). White squares indicate origin of inset. Scale bar, 20 µm. **f**, Time course of H4 cells transduced with AAV overexpressing TDP-43venus after infection with HSV-1 (MOI 0.3) and treatment with Acyclovir (ACV) 4 h post-infection. TDP-43venus particle counts were normalized to the count of the first measurement and to the fold of the corresponding time points of the control (mock). Dots represent the mean±SEM, n = 3, one biological replicate. **g**, Quantification of nuclear TDP-43venus foci area of H4 cells co-expressing respective ICPs in (**h**) using Foci Analyzer. Lines represent mean±SEM. Dots represent individual cells, n = 17-94, three independent biological replicates, ***p<0.001 Kruskal-Wallis test followed by Dunn’s multiple comparisons test. **h**, Representative immunofluorescence images of H4 cells co-transfected with TDP-43venus (green) and indicated HSV-1 immediate-early gene (ICP0, ICP4, ICP27, ICP22myc, ICP47myc) (red). Merged images show additionally DAPI (blue). Scale bar, 10 µm. **i**, Representative fluorescence images of TDP-43venus in H4 cells transduced with AAV-TDP-43venus for 3 days and subsequently infected with HSV-1 and HSV-1 ΔICP0 (MOI 0.3) at 0 h and 16 h post-infection (p.i.). Scale bar, 15 µm. **j**, Quantification of TDP-43venus particle in (**i**) normalized to the count of the first measurement and to the fold of the corresponding time points of the control (mock). Dots represent the mean±SEM, n = 9, three independent biological replicates. A linear regression model with time and treatment as fixed effects demonstrates a significant effect for treatment (***p<0.001) and treatment:timepoint interaction (***p<0.001). Pairwise comparison with estimated marginal trend contrasts compared HSV-1 and HSV-1ΔICP0, demonstrating a significant difference (***p<0.001).

These results suggest that expression of the immediate early HSV-1 protein ICP0 promotes delocalization of TDP-43.

### Disruption of PML-NBs by HSV ICP0 prevents stress-induced SUMOylation of TDP-43

HSV-1 ICP0 and its homologs in other herpesviruses are multifunctional proteins: ICP0 acts as an E3 ubiquitin ligase to target PML-NBs^31^, via its RNF8 binding it controls histone ubiquitination^32^, the cyclin D3 interaction regulates the cell cycle^33^ and binding to CoREST was reported to disrupt the histone deacetylase repressor complex^34^. To understand which ICP0 function is required to delocalize TDP-43, we generated ICP0 mutants selectively disrupting the E3 ubiquitin ligase activity (C116G/C156A), the RNF8-binding site (T67A), the cyclin D3–binding site (D199A), or the CoREST-interaction motif (D671A/E673A)^35–37^ (Fig. 5a). The overall mean fluorescence intensity of TDP-43venus was not affected in any of the ICP0 mutants (Extended Data Fig. 5a). Expression of wild-type (WT) ICP0, as well as the T67A, D199A, resulted in robust formation of nuclear TDP-43 puncta (Fig. 5b, c). D671A/E673A mutation reduced the number of puncta slightly, but still increased the number of TDP-43 puncta significantly compared to the vector control. Strikingly, the mutation of C116G/C156A resulted in a loss of TDP-43 delocalisation, suggesting that the E3 ligase activity modulated TDP-43 localisation directly or indirectly. The primary role of its E3 ubiquitin ligase function is to dismantle the anti-viral PML-NBs, thereby promoting efficient viral gene expression and productive infection^38^. As expected, upon ICP0 expression, the typical nuclear puncta of PML-NBs were lost^39,40^ (Fig. 5b). Intriguingly, upon overexpression of the ICP0 mutant proteins, the average number of PML-NBs significantly negatively correlated with the average size of TDP-43 puncta (r=-0.79, p=0.03), suggesting that the ability of the different ICP0 proteins to disrupt PML-NBs was correlated with TDP-43 delocalization (Extended Data Fig. 5b). In line with this, single-cell correlation analyses revealed that the number of PML bodies negatively correlated (r=-054, p<0.001) with the nuclear TDP-43 foci area, in both vector and ICP0 expressing H4 cells (Fig. 5d). ICP0-mediated ubiquitination of PML causes its proteasomal degradation^41^. To substantiate that the PML-NB disruption function of ICP0 was responsible for delocalization of TDP-43, we inhibited proteasome activity using MG132. Proteasome inhibition largely retained PML nuclear bodies upon infection with HSV-1 and was accompanied by a pronounced reduction in HSV-1–induced nuclear TDP-43 foci (Fig. 5e, f). Again, the overall fluorescence signal of TDP-43 was unaffected (Extended Data Fig. 5c). PML bodies are not only targeted by HSV-1 ICP0, but a similar mechanism is employed by HSV-2 ICP0. Homologous herpesviral E3 ligases, such as HCMV IE1, VZV ORF61, EBV BNRF1, or KSHV ORF75, may disrupt PML-NB integrity but do not target PML for degradation. None of the other herpesviral HSV-1/2 ICP0 homologues delocalize TDP-43 significantly (Extended Data Fig. 5c, e)^42–44^. SUMOylation of TDP-43 with SUMO2/3 in PML-PBs promotes TDP-43 solubility upon stress^5,6^. Thus, we hypothesized that PML disruption by ICP0 may impact TDP-43 SUMOylation. To this end, FLAG-tagged TDP-43 was purified from vector or ICP0 expressing cells and the SUMOylation status was analyzed. In vector control cells, SUMO2-modified TDP-43 was readily detected following arsenite treatment that was used to induce cellular stress^45^. In contrast, HSV-1 ICP0 expression led to a marked reduction in SUMO2-modified TDP-43 by over 80% (Fig. 5g, h). Mono-SUMOylation was even more drastically reduced (Extended Data Fig. 5f).

**Fig. 5:**
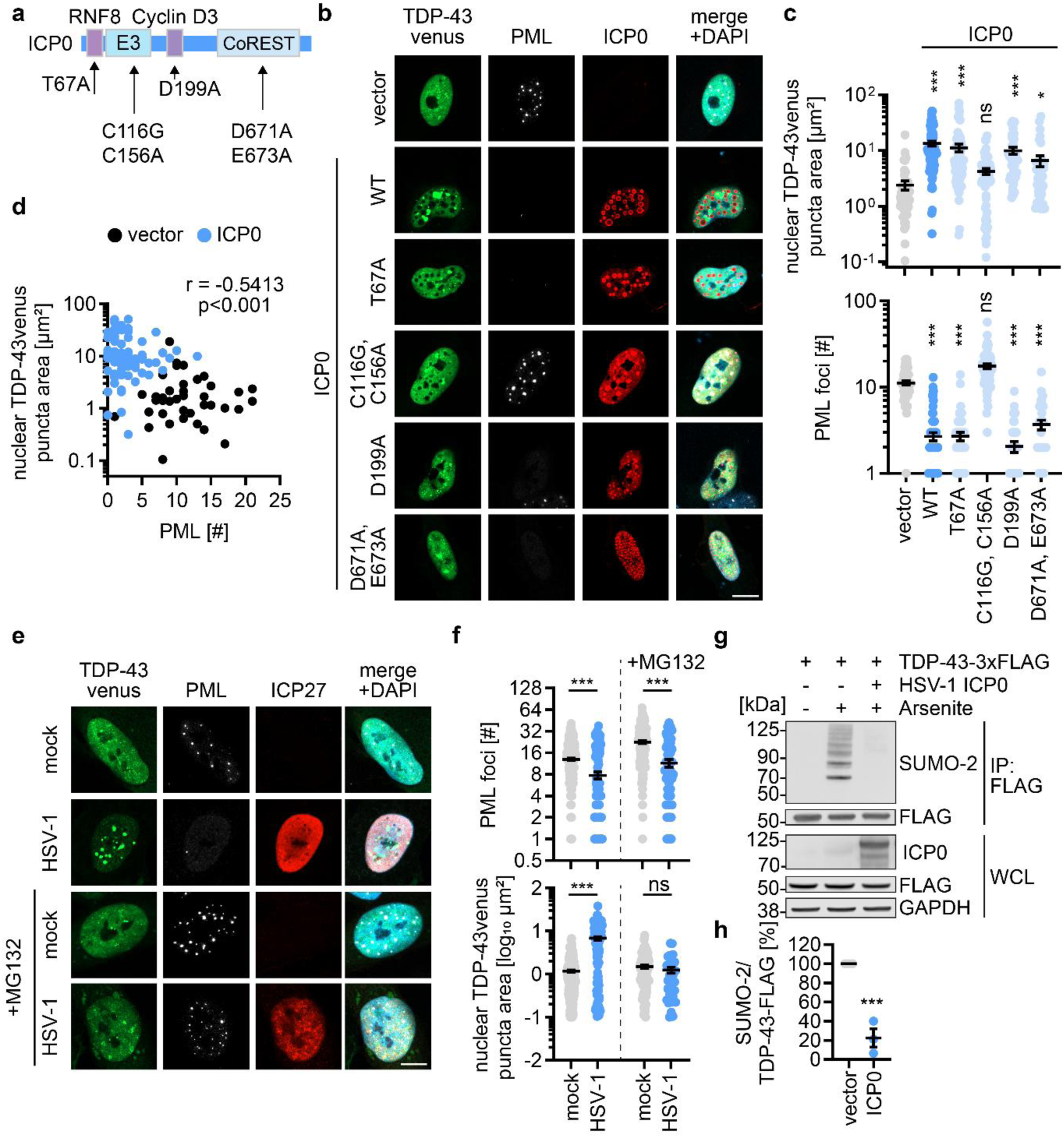
PML body disturbance by ICP0 is responsible for TDP-43 delocalization and loss of SUMOylation. **a**, Schematic overview of the HSV-1 ICP0 gene with mutations to disrupt either RNF8 binding site (T67A), E3 ligase function (C116G/C156A), Cyclin D3 binding site (D199A) or CoREST interaction (D671A/E673A). **b**, Representative immunofluorescence images of H4 cells co-transfected with TDP-43venus (green) and either vector, wild-type (WT) ICP0 or indicated ICP0 mutant. Cells were fixed 24 h post-transfection and stained for PML (white), ICP0 (red) and DAPI (blue). Scale bar, 10 µm. **c**, Quantification of nuclear TDP-43venus foci area (µm^2^) (top) and PML foci count (bottom) of co-transfected cells in (b) using Foci Analyzer (ImageJ). Lines represent mean±SEM, dots represent individual cells (n = 37-74) of three independent biological replicates. *p<0.05, ***p<0.001 Kruskal-Wallis test followed by Dunn’s multiple comparisons test. **d**, Scatter plot of total nuclear TDP-43venus foci size (y-axis) against the nuclear PML dot count (x-axis). Each dot represents one corresponding measurement from (c). Blue dots represent cells overexpressing ICP0 WT, black dots represent cells transfected with vector. For association testing, all events were taken together in Spearman’s rank correlation test (r = –0.54, ***p<0.001, n = 118). **e**, Representative immunofluorescence images of H4 cells transduced with lentiviruses encoding TDP-43venus (green). After 3 days, cells were infected with HSV-1 and 1 h post-infection treated with proteasome inhibitor MG132 or DMSO, respectively. Cells were fixed 17 h post-infection and stained for PML (white), ICP27 (red) and DAPI (blue). Scale bar, 10 µm. **f**, Quantification of PML foci (top) and total nuclear TDP-43venus foci area (µm^2^) (bottom) in (e). Lines represent mean±SEM. Dots represent individual cells, n = 58-362, three independent biological replicates, ***p<0.001 Kruskal-Wallis test followed by Dunn’s multiple comparisons test. **g**, Representative immunoblots showing SUMOylation status of FLAG-tagged TDP-43 purified by anti-FLAG immunoprecipitation (IP) from whole cell lysates (WCL) of HEK293T cells at 40 h post-transfection. Cells were co-transfected with HSV-1 ICP0 and/or stimulated with arsenite (500 µM, 1 h), as indicated. IP and WCL was analyzed by immunoblotting. Blots were stained with anti-SUMO2, anti-FLAG, anti-HSV-1 ICP0 and anti-GAPDH. **h**, Quantification of the full SUMO-2 lane in the TDP-43 IP of (**g**) normalized to the vector control. n = 3, **p<0.01 paired t test.

These findings demonstrate that disruption of PML by HSV ICP0 E3 ligase activity reduces TDP-43 SUMOylation and thereby influences its solubility.

## DISCUSSION

Emerging evidence links viral infections and neurodegenerative diseases^11,13–15^. However, the precise associations and underlying molecular mechanisms have been unclear. In this study, we identified a previously unrecognized association of HSV infection and ALS in independent population data and ALS patient cohorts. Furthermore, HSV-1 infection induced key molecular hallmarks of ALS in cell lines, primary neurons, human iPSC–derived motoneurons and forebrain organoids, as well as in mouse models. Our mechanistic analyses support a model in which the HSV-1 ICP0 E3 ligase disturbs PML-NB integrity, resulting in a loss of TDP-43 SUMOylation, which eventually causes its delocalization, hyperphosphorylation and insolubility.

Different viruses, most prominently certain herpesviruses, have previously been implicated in the development of NDDs^15^. Previous population data had linked HSV-1 primarily to dementias, typically AD^22,46,47^. Mechanistic analyses suggest that HSV-1 promotes signs of AD pathology including Aβ accumulation, tau phosphorylation, neuroinflammation, oxidative stress, and memory impairment in mice^23,48^. More recently it was shown that HSV-1 infection promotes the release of phosphorylated tau via extracellular vesicles *in vitro* and *in vivo*^49^. Our study now adds ALS to the spectrum of NDDs linked to diseases associated with HSV. While future studies assessing clinical relevance based on our serology and epidemiological data are clearly needed, the notion that specifically HSV-1 and 2 may be important in ALS pathogenesis is corroborated by our mechanistic data showing that only HSV-1/2 drive TDP-43 delocalization, but no other herpesviruses tested. However, our data do not exclude the possibility that other viruses may be associated with ALS as well. Cytosolic and nuclear aggregates of TDP-43 are not only associated with the pathology of ALS but also Frontotemporal dementia (FTD) and a more recently recognized age-associated dementia termed limbic-predominant age-related TDP-43 encephalopathy^50^. Thus, the impact of HSV-1 on TDP-43 may have implications for other NDDs. While this study mainly focuses on sporadic ALS, which comprises ∼90% of all ALS cases, genetic risk factors of ALS may also increase the likelihood or severity of HSV-1 reactivation or initial infection since many ALS-associated genes (e.g. OPTN, TBK1)^51^ contribute to the innate immune defense against viruses, including HSV-1^52–54^.

Mechanistic analyses revealed that disruption of PML-NBs by HSV-1/2 ICP0 drives TDP-43 delocalisation. PML-NBs are membrane-less nuclear, multiprotein regulatory hubs that are important for post-translational modifications, nuclear protein quality and anti-viral defenses^55^. Thus, many viruses, including HSV, disrupt or modify PML nuclear bodies^56^. For example, ICP0 of HSV-1 promotes proteasomal degradation of PML itself and the PML-NB components IFI16 and HIRA during the early phases of infection^38^. Other herpesviruses employ different strategies: IE1 of HCMV disrupts PML-NBs, but does not lead to PML degradation; VZV ORF61 disturbs PML-NBs but also does not cause PML degradation^42,57^. Since we observed TDP-43 delocalisation only with HSV ICP0, and not ICP0 homologous of other human herpesviruses that do not target the protein PML directly, we infer that degradation of PML itself may promote TDP-43 delocalisation. Recently, it was demonstrated that SUMO2/3 conjugation of TDP-43 in PML-NBs prevents its aggregation and regulates its subcellular localization^6,58,59^. Our data now shows that HSV-1 ICP0 prevents efficient SUMO2/3-conjugation to TDP-43 under cellular stress, thereby likely mediating its delocalisation and oligomerisation.

Interestingly, a loss of PML NBs in post-mortem brain tissue from familial ALS/FTD has been observed^60^. In addition, aberrant chromatin accessibility or alternative polyadenylation (APA) were suggested to be additional hallmarks of ALS pathology^61^. For example, our previous work showed that ALS patient samples exhibited a more compact chromatin state compared to healthy controls^61,62^. APA changes were also observed in ALS/FTD-related post-mortem human tissue and linked to TDP-43 pathology^63^. It is interesting to note, just like degradation of PML-NBs, major remodeling of chromatin accessibility as well as massive changes in host mRNA processing including increased APA occur during HSV-1 infection^64,65^. Whether molecular hallmarks of ALS are partially signatures of previous HSV-1 replication/reactivation events warrants future studies.

HSV-1 seroprevalence in the general adult population is around 65%^66,67^ prompting the questions as to why ALS is a very rare disease (6 per 100 000)^68^. This suggests that either HSV-1 infection alone cannot be causal for ALS and additional factors like genetic risks or stress are required, or that the infection event driving pathology is rare or needs to occur multiple times to cross a threshold to cause self-propagation of TDP-43 aggregates. It is tempting to speculate that subclinical, albeit rare mild or self-limiting reactivation of HSV-1 into the CNS drives initial TDP-43 delocalization.

Our data associate TDP-43 pathology on a molecular level to HSV-1 infected cells. ALS is clinically mainly defined by degeneration of motoneurons^4^. However, HSV-1 classically infects sensory neurons and establishes latency in the trigeminus ganglia^29^. Thus, to cause ALS pathology, HSV-1 would need to spread from sensory neurons to motoneurons, or TDP-43 spreading from HSV-1 infected neurons. However, a few studies arguing that HSV-1 might not be completely absent from motoneurons, e.g. in rats, a transneuronal transfer of HSV-1 from sensory neurons to motoneurons was reported^69^. In addition, in latency mouse models, HSV-1 was found in the motoneuron containing Pr5 region^70^ and using a gene therapy vector, HSV-1 was able to deliver its cargo to motoneurons *in vivo*^71^. In human herpes simplex virus encephalitis cases, HSV-1 was detected in motoneurons^72^. Finally, motoneurons may be rarely infected, but might be more vulnerable to HSV-1 induced TDP-43 pathology. Motoneurons express lower levels of IFNAR1/2 (type I interferon receptors) and STAT1, which are essential for canonical interferon signaling^73^ that regulates PML expression^74^. PML is important for suppression of lytic HSV-1 gene expression^38^. Once PML-mediated chromatin control is weakened, HSV replication resumes. It is tempting to speculate that HSV-1 exploits these differences between sensory and motoneurons and reactivates more efficiently in motoneurons. Of note, an altered interferon response has been reported in ALS^75,76^.

The association between HSV-1 and ALS may inspire innovations in ALS therapy and prophylaxis^14^. Recently, large population studies demonstrated that vaccination against herpes zoster, a debilitating skin disease caused by reactivation of latent VZV, was associated with a lower risk of dementia^11,13,77^. Current antiviral drugs targeting HSV-1 have shown benefits in observational studies of dementia risk^14,15^, suggesting that viral suppression may modify neurodegenerative trajectories. It is thus tempting to hope that new developments in preventing HSV-1 infection or reactivation by a vaccine or therapeutic intervention with HSV-1 may reduce disease burden of ALS.

## MATERIALS AND METHODS

### Study cohorts and ethical approval

Sporadic ALS (sALS) patients and as well as healthy controls were recruited at the university clinic Ulm, Hannover Medical School, Department of Neurology, University Medical Center Göttingen and Department of Neurology Halle-Wittenberg and Paracelsus-Elena Clinic Kassel. All human experiments were performed in accordance with the declaration of Helsinki and approved by the Ethics Committee of Ulm University and the respective medical centers. Informed consent was obtained from all participants included in the study. All patients were diagnosed according to the El-Escorial criteria for definite ALS. Patients were considered sporadic based upon a negative family history. Individuals with confounding conditions affecting the immune system were excluded. Healthy controls were chosen to match the patient cohorts or mutation carrier cohort, respectively, regarding age and sex.

FFPE liver tissue samples from patients with clinically and diagnostically confirmed hepatitis caused by HSV-1, hepatitis B virus (HBV), or hepatitis C virus (HCV) were obtained for diagnostic purposes and provided by the BioBanks of Erasmus MC. In accordance with the institutional “Opt-Out” system defined in the Dutch National Code of Good Conduct (Code Goed Gebruik, May 2011), surplus diagnostic tissue was made available for research use in this study.

### Serum isolation

Whole venous blood was collected from all participants with a standard S-Monovette serum drawing system (Sarstedt). Blood samples were processed within 4 hours after blood collection. The samples were centrifuged at 2,000 x g for 10 min at 4 °C. The upper liquid phase was transferred into a new tube and stored at –80 °C.

### Analysis of electronic healthcare claims data

The electronic healthcare claims data consists of a random sample of 2.2% (n = 250,000) of the total insured population of German healthcare provider selected at the beginning of 2004 and followed up until the end of 2019. We used data of 238,440 individuals aged 50 and older who had no diagnosis of ALS in their medical history during the year 2004. We identified 113 patients with a valid new diagnosis of ALS between the first quarter of 2005 and the third quarter of 2019. We used a multi-stage internal validation process to minimize the problem of false-positive cases. We used only verified diagnoses from the outpatient sector or only discharge or secondary diagnoses from the inpatient sector. An ALS diagnosis must be confirmed by at least one second diagnosis in a subsequent quarter, or the patient must have received an outpatient and an inpatient diagnosis in the same quarter. We defined only ALS diagnoses as valid if at least one third of all quarters after the initial diagnosis were given an ALS diagnosis, as it is a very care-intensive disease. We included only patients with at least one prescription for riluzole and, we excluded all patients with at least one diagnosis of ALS in 2004, even if the diagnosis was not confirmed. Any new case meeting the above criteria between 2005 and 2019 was considered an incident ALS case. The date of the first diagnosis is subsequently referred to as the diagnosis date. Each of the 113 new ALS cases was matched to four controls by age (in ten-year age groups), gender, and time since observation using incidence density sampling (Stata command sttocc). For the controls, the index date was set to the date of ALS diagnosis of the matched cases. We adjusted the regression model for prescription of antivirals for systemic use (ATC code: J05AB) and antivirals for dermatological use (ATC code: D06BB) and for the weighted Charlson comorbidity index, according to Quan et al. 2011^78^. All analyses were performed with Stata 16.1 MP (StataCorp LLC).

### Herpesvirus serology

The following commercial tests were used to determine the antibody titers from patient sera according to the manufacturer’s instructions. CMV IgG was quantified using the Liaison CMV IgGII and VZV IgG using the Liaison VZV IgG HT on a Liaison XL instrument (Diasorin). EBV VCA IgG was quantified using the EBV VCA IgG assay on an Architect 1000 instrument (Abbott). HSV IgG was quantified using the Herpes Simplex Virus 1/2 IgG assay on an Immunomat instrument (Virion-Serion). The quantity of some serum samples was limited; therefore, samples were diluted 1/10 in PBS to obtain comparable results. Only antibody titers with unequivocal interpretation were included, and only samples for each all 4 antibody titers were available. The multiple logistic regression model was fitted with the ‘glm()’function from R (v.4.5.2) ‘stats’ package (v4.5.2) with “ALS symptomatic” as dependent variable and centered age, sex, CMV, HSV-1, VZV and EBV status as dependent variables and “bionomial” test family.

### Cell culture and media

Human H4 neuroglioma cells (HTB-148, American type culture collection (ATCC)), HEK293T (CRL-3216, ATCC), HEK-293 (CRL-1573, ATCC), Lenti-X 293T (VP001, Merck), Vero E6 (CRL-1586, ATCC) and MRC-5 fibroblasts, U2OS (kindly provided by Robert Tjian and Xavier Darzacq (UC Berkeley)) were cultured in Dulbecco’s Modified Eagle Medium (DMEM, Gibco) supplemented with 10% (v/v) fetal bovine serum (FBS, Gibco) and 2 mM L-glutamine (PAN-Biotech) (hence DMEM(++)). Cells were cultivated in a 5% CO_2_ humidified incubator at 37 °C and regularly tested for mycoplasma contamination. Cells were only used if negative.

### Primary neurons

Culture plates were coated with poly-L-lysine (PLL, 0.1 mg/ml) to support adherent growth of primary murine neurons. Plates were incubated for at least 4 h at 37 °C, washed with PBS, and stored in PBS+ (with Ca²⁺/Mg²⁺) at 4 °C. On the day of neuron isolation, PBS+ was replaced with DMEM(++) (with addition of 1% Penicillin/Streptomycin (P/S)), and plates were equilibrated at 37 °C and 5% CO₂. Primary cortical neurons were isolated from C57BL/6J mice at postnatal day 0–1 under approved guidelines of the Animal Research Center, Ulm University. Pups were decapitated, and brains were removed into ice-cold HBSS. Hemispheres were separated, the cortex was dissected, meninges were removed, and tissue was collected in cold HBSS. Tissue was washed three times, and 1–2 ml HBSS were retained before adding 250 µl trypsin (2.5%, no EDTA). The suspension was incubated for 20 min at 37 °C and 5% CO₂ with gentle inversion every 5 min. DNase (100 µl, 0.1 mg/ml) was added and incubated for 5 min at room temperature. The supernatant was removed, trypsinization was stopped with 5 ml DMEM(++) (+1% P/S), and cells were washed twice with HBSS, followed by an additional 100 µl DNase. After two washes with DMEM(++), cells were homogenized in 2.5 ml DMEM(++), supplemented with 5 ml DMEM(++), and passed through a 100 µm strainer. The strainer was rinsed twice with 10 ml DMEM(++), and the suspension was centrifuged at 300 × g for 5 min. The pellet was resuspended in 2–3 ml neurobasal medium+++ (2% B27, 1% P/S, 1% L-glutamine). Cells were counted using a Neubauer chamber and plated in neurobasal+++ in PLL-coated 24-wells dishes (0.15 × 10⁶ cells in 500 µl per well). Cultures were maintained at 37 °C and 5% CO₂. After 2 days, half of the medium was replaced with fresh neurobasal+++. On the day of transduction, one-fifth of the medium was exchanged with double the volume of fresh medium. AAV TDP-43venus vectors (Titre 2.94×10^10^ gc/ml; 5.1 µl) were then added, and cultures were incubated for 3 days. Cells were washed three times with PBS and infected with HSV-1 GFP with an approximate multiplicity of infection (MOI) of 0.3. After 17 h, cells were then fixed using 4% PFA for 10 min and kept in PBS at 4 °C.

### hiPSC culture and differentiation into forebrain organoids

The hiPSC line SCTi003-A and SCTi004-A were purchased from StemCell Technologies and maintained at 37 °C and 5% CO_2_. Cells were cultured in mTeSR Plus (StemCell Technologies #100-0276) on 6 well plates (Sarstedt) coated in Matrigel (Corning, #354277) diluted in DMEM-F12 (Gibco, #31331-028) according to manufacturer’s instructions and passaged with Dispase (StemCell Technologies #07913) when confluency reached 60–70%. Cells were cryopreserved as colonies in mFreSR (StemCell Technologies #05855). Forebrain organoids were differentiated from hiPSC using a previously published protocol with minor modifications^79^. Briefly, at day 0 hiPSCs were dissociated to single cells using Accutase solution (Millipore, #SCR005) and seeded 15,000 cells/well in 96-well ultralow attachment plate for the formation of embryobodies in neural induction medium (NIM) with 10µM ROCKi (Abcam, #Ab120129). NIM is composed of: DMEM-F12 (Gibco, #31331-028), 20% KnockOut (Gibco, 10828028) serum replacement, 1% NEAA (Gibco, #11140-035), 1% β-mercaptoethanol (Millipore, #ES-007-E), 1% antibiotic-antimycotic (Sigma Aldrich #A5955), 10 µM SB-431542 (StemCell Technologies, #72234), 2 µM dorsomorphin (StemCell Technologies, #72102), 1% Neurocult supplement without vitamin A (Stemcell Technologies, 05731), 0.5% N2 supplement (Gibco, 17502-284). Half media change was performed daily for 7 days without ROCKi. At day 8, organoids were transferred to 24-well ultralow attachment plates and media was replaced with neural expansion medium: Neurobasal-A Medium (Gibco, #21103049, 2% Neurocult supplement without vitamin A, 1% GlutaMAX supplement (Gibco, #35050061), 20 ng/ml EGF (Peprotech, #AF-100-15) and 20 ng/ml FGF2 (Peprotech, #100-18B). 500 µl of NEM was added daily until day 15 and every other day until day 24. On day 25, the medium was replaced with neural differentiation medium: Neurobasal-A Medium (Gibco, #21103049, 2% Neurocult supplement without vitamin A, 1% GlutaMAX supplement (Gibco, #35050061), 20 ng/ml BDNF (Peprotech, # 450-02), 20 ng/ml NT3 (Peprotech, # 450-03) 200 µM Ascorbic Acid (PanReac AppliChem, #131013.1208), 50 µM of cAMP (Enzo, #BML-CN125-0030) and 10 µM of DHA (Sigma Aldrich, #D2534). Full media change was performed twice a week. On day 47, forebrain organoids were maintained in neural maturation medium: Neurobasal-A Medium (Gibco, #21103049, 2% Neurocult supplement (StemCell Technologies, 05711). Full media change was performed twice a week.

### Mice

C57BL/6J mice were purchased commercially (Jackson laboratories, stock #000664) and housed in a specific pathogen-free barrier facility (RT of 20-26 °C and humidity of 30-70%) with a 12 h dark/light cycle and unlimited access to standard food and water at the Cleveland Clinic Florida Research and Innovation Center (CC-FRIC). All mice used in this study were handled in accordance with policies and procedures that were approved by the Institutional Animal Care and Use Committee (IACUC) at CC-FRIC.

### Virus stock production

#### HSV-1, HSV-2

HSV-1 (Strain 17+)^80^, HSV-1 GFP (Strain F)^81^, and HSV-2 (Clinical isolate, kindly provided by Sebastian Michel, Institute of Virology, Ulm University Medical Center) were amplified using Vero E6 cells (ATCC CRL-1586). 5 x 10^6^ Vero cells were seeded in DMEM(++) in a T175 flask and infected with respective virus the day after with an MOI of 0.01. Cells were daily checked for cytopathic effects (CPE) and harvested after 70-80% of cells showed a CPE. Cells were detached by scratching into the supernatant (SN). Sonification (10 min) of the cells was applied and afterwards, cell debris was removed via centrifugation (10 min, 4 °C, 750 g). Aliquots were stored at –80 °C. Virus titres for all viruses were expressed as the 50% cell culture infective dose (TCID₅₀), calculated according to the method of Reed and Muench^82^. In brief, virus dilution series was added eightfold to cell type of interest. At day 3 post-infection, virus positivity was assessed by CPE.

BACs of the HSV-1 F strain ΔICP0 mutant and the corresponding F wild-type strain were a gift from B. Roizman (University of Chicago)^83^. BAC DNA was prepared from a monoclonal transformant, sequence-verified by long-read sequencing (Plasmidsaurus), and transfected into U2OS cells using Lipofectamine 2000 (Thermo Fisher). Virus was propagated, harvested, and stored as previously described^84^. Viral progeny in cell-free supernatants were titrated on U2OS cells by determining the TCID₅₀ using eight replicates per dilution. Infectious units (FFU/ml) were calculated using the Spearman–Kärber method^85^.

HSV-1 (strain KOS, kindly provided by D. Knipe at Harvard University) was propagated in African green monkey kidney epithelial cells (Vero, CCL-81; ATCC) and titered in the same cell line by plaque assay as previously described^86^.

#### MeV

For MeV (vac2 strain^87^, kindly provided by Karl-Klaus Conzelmann, Max von Pettenkofer-Institute, LMU, Munich) propagation, confluent Vero E6 cells of a T175 flask were trypsinated and pelleted at 1600 rpm for 3 min at 4 °C. Afterwards, cell pellet was resuspended in MeV, reaching an MOI of 0.01. Infected cells were incubated for 1 h at 37 °C with gentle shaking every 10-15 min. Afterwards, cells were diluted with respective medium and seeded into a T175 flask, followed by incubation at 37 °C until syncytia formation was detected. At this point, medium was replaced by 5 ml Opti-MEM and cells were harvested using a cell scraper. Cell suspension was sonicated for 10 min to disrupt and to release the virions into the supernatant. Afterwards, cell debris was pelleted (1600 rpm, 3 min, 4 °C) and the supernatant was stored at –80 °C.

#### EMCV

EMCV (EM strain, ATCC) was propagated on HEK293T cells as previously described^88^. Briefly, 18 h post-infection, the supernatant was cleared of cell debris by centrifugation (1600 rpm, 3 min, 4 °C) and the supernatant was stored at –80 °C. Virus titres were determined as the TCID₅₀, calculated according to the method of Reed and Muench^89^.

#### HCMV

The HCMV, TB40/E strain was propagated in human foreskin fibroblast (HFF)^90^. After infection, the supernatant was cleared of cell debris via centrifugation, aliquoted and stored at –80 °C. Titration was performed on HFF by staining for IE1. IE1 positive cells were counted via ImageXpress Pico (Molecular Devices), giving viral titers expressed as IE protein-forming units (IEU).

#### VZV

For propagation of cell-free VZV, MRC-5 fibroblasts were seeded in a T175 flask, cultured in Dulbecco’s Modified Eagle Medium (DMEM, Gibco) supplemented with 2% (v/v) fetal bovine serum (FBS, Gibco) and 2 mM L-glutamine (PAN-Biotech). At 80% confluency, cells were infected with 100 µl Human herpesvirus 3 VR-1367 (ATCC). When cells started to show cytopathic effect (CPE), infected cells were mixed with uninfected MRC-5 at a ratio of 1 to 6 and seeded in 15 cm dishes. For preparation of cell-free VZV stocks, PSGC buffer was prepared as follows: 5% sucrose (w/v) (Sigma-Aldrich), 0.1% (w/v) L-glutamic acid monosodium salt hydrate (Sigma-Aldrich) and 10% (v/v) FBS (Gibco) in 1x PBS, sterile-filtered. Virus was harvested when CPE reached 30%. Cells were washed once with cold 1x PBS, scraped into 6.5 ml cold PSGC buffer per dish and sonicated for 10 min. Cell debris was removed via centrifugation (15 min, 4°C, 1000 g). Supernatants were transferred to 100 kDa Amicon centrifugation filters (Merck) and 20x concentrated via centrifugation (3300 g, 4°C). Cell-free VZV was stored at –80°C and titrated on H4 cells.

#### AAV-TDP-43venus

To produce AAV-TDP-43venus, 293T cells were split 1 to 3 on 20 15 cm standard cell culture dishes (Sarstedt) 18-20 h before transfection in order to reach ∼60-70% confluency at the time of transfection. Directly before transfection the medium was exchanged from 25 ml DMEM/ 10% FCS/ 1X GlutaMAX to 15 ml DMEM/ 1% FCS/ 1X GlutaMAX (Gibco, Thermo Fisher Scientific) per plate. Cells were transfected with 35 µg total DNA/plate, complexed with PEIpro (Polyplus, Sartorius) at a ratio of 1:2. The pAAV_TDP-43 venus transfer plasmid, the adenoviral pHelper plasmid (Cell Biolabs) and the AAV helper plasmid pRep2-Cap6.2 (kindly provided by Dr. Benjamin Strobel, Boehringer Ingelheim Pharma GmbH & Co. KG^91^) were used in an equimolar ratio. At 3 h post-transfection 10 ml of DMEM/ 11% FCS/ 1X GlutaMAX were added in order to have a final volume of 25 ml with an FCS content of 5%. Cells were incubated at 37 °C, 5% CO_2_ and 95% humidity for 72 hours. Then they were scraped off and centrifuged together with the virus-containing medium at 400 g at 4 °C for 15 min. The virus-containing medium was filtered through a 0.45 µm PES membrane and AAVs were precipitated by addition of 40% PEG-8000 (sterile-filtered) to a final percentage of 8% overnight at 4 °C. Production cells were resuspended in lysis buffer (50 mM Tris, 10 mM MgCl_2_, 0.001% Pluronic F-68, 150 mM NaCl, pH 8; sterile-filtered, 5% of culture volume) and underwent three freeze-thaw cycles (liquid N2, 37 °C water bath). 50 U of Denarase (c-LEcta) were added per ml lysis buffer and nucleic acids were digested for 45 min at 37 °C. Then, Denarase was inhibited by addition of EDTA to a final concentration of 5 mM, and cell debris was spun down at 2500 g and 4 °C for 15 min. The virus-containing lysate was transferred to a fresh Falcon tube and AAVs were precipitated with 8% PEG-8000 at 4 °C overnight. After 30 min of centrifugation at 2500 g and 4 °C precipitated AAVs from the virus-containing medium and the cell lysate were resolved in 13 ml resolution buffer (1 M NaCl, 50 mM Tris, 0.001% Pluronic F-68, pH 8.0, sterile-filtered) overnight on a rotating platform at 4 °C. After a final centrifugation (2500 g, 4 °C, 30 min), the AAV-containing suspensions were transferred to a 39 ml Quick-Seal Ultra-Clear Tube (Beckman). An iodixanol gradient was set up by underlaying the AAVs with 9 ml of 15% OptiPrep (iodixanol) in PBS-MKN (DPBS with 1 mM MgCl2, 2.5 mM KCl and 1 M NaCl, sterile filtered), 6 ml of 25% iodixanol in PBS-MK (DPBS with 1 mM MgCl2, 2.5 mM KCl, sterile filtered), 5 ml of 40% iodixanol and 5 ml of 54% iodixanol. Tubes were filled up with resuspension buffer, sealed and centrifuged for 2 h at 200,000 g and 18 °C in an Optima XE-90 Ultracentrifuge (70 Ti fixed-angle rotor, Beckman Coulter). The AAV-containing 40% layer was drawn with a syringe and concentrated by ultrafiltration (Amicon Ultra-15, 100 kDa MWCO). Thereby the iodixanol/PBS-MK was exchanged to formulation buffer (2.5 mM KCl, 1 mM MgCl2, 0.001% Pluronic F-68 in PBS, pH 7.4, 10% glycerol). The genome-copy titer of purified vector stock was determined by qPCR using the AAVpro Titration Kit (TaKaRa). Purity on the protein level was confirmed by silver staining after polyacrylamide gel electrophoresis^92^.

#### Lentivirus

For lentivirus production, 2.5 x 10^6^ Lenti-X 293T cells per dish were seeded in six 100 mm dishes each with DMEM(++) and co-transfected with 45 µg of the pLenti plasmids (TDP-43venus, TDP-43mCherry and mCherry), the LVTetO plasmid TDP-43halo^93^ containing the gene of interests (GOI) and two packaging plasmids psPax2 (45 µg) and pMD2.G (18 µg) per dish, respectively. Cells were transfected using calcium-phosphate precipitation. Reagents were incubated for 30 min at RT, after addition of 180 µl 2.5 M CaCl_2_, 3000 µl 2x HBS and 109.4 µl ddH_2_O. Transfection mixture (560 µl) was then added to each cell plate and substituted with culture medium after 6 h. Lentiviruses were isolated 2 days after transfection. Therefore, medium of all dishes was collected in 50 ml tubes and filtrated using a 0.2 µm sterile filter. Filtrated supernatant was transferred to two ultracentrifuge tubes per condition and underlayered with 3 ml of 20% sucrose. To isolate viral particles, ultracentrifuge tubes were centrifuged at 100,000 x g for 2 h at 4 °C (Centrifuge Beckmann Coulter Optima XPN-80, rotor type 70). Supernatant was discarded carefully and pellets were dissolved with 100 µl PBS per tube. Flow cytometry was used for validation and titration of each Lentivirus stock.

#### Expression constructs

The generation of the pcDNA3-TDP-43-L1, pcDNA3-TDP-43-L2 and pcDNA3-TDP-43venus plasmids has been previously described^91^, as has the LVTetO lentiviral TDP-43halo construct^93^. The pAAV_TDP-43venus construct was generated by cloning an insert containing XhoI, NotI restriction sites into the transfer plasmid pAAV, using the primers AAV_TDP_XhoI_fw and AAV_venus_NotI_rev (Table 1) and the TDP-43venus plasmid^26^ as template. For the pcDNA3-3xFLAG-TDP-43 construct, an insert harbouring BamHI and XbaI restriction sites was produced using the primers 3xFLAG_TDP-43_BamHI_fw and 3xFLAG_TDP-43_XbaI_rev (Table 1), with the TDP-43venus plasmid^26^ serving as template. The pLenti-TDP-43venus plasmid was cloned by generating an insert flanked by PacI and SpeI restriction sites using the primers pLenti_TDP-43_PacI_fw and pLenti_TDP-43venus_SpeI_rev (Table 1). Construction of the pLenti-TDP-43mCherry plasmid required two separate inserts: a TDP-43 fragment containing PacI and BamHI restriction sites, amplified with primers pLenti_PacI_TDP_fw and pLenti_BamHI_TDP_rev (Table 1), and an mCherry fragment containing BamHI and SpeI restriction sites, generated using the primers pLenti_BamHI_mCherry_fw and pLenti_SpeI_mCherry_rev (Table 1) and theCRY2olig-mCherry plasmid (addgene, #60032) as template. The HSV-1 proteins ICP0, ICP4, ICP22, ICP27, ICP47 (Strain 17, NCBI reference sequence: NC_001806.2) were codon optimized and synthesized in pTwist EF1 Alpha vectors by Twist Bioscience. The pcDNA3-MycICP22 and pcDNA3-ICP47Myc plasmids were cloned by creating inserts containing BamHI and EcoRI restrictions sites; primers are listed in Table 1. ICP0 mutants (T67A, D199A, E671A E673A, C116G, C156A) were constructed using the Q5 Site-Directed Mutagenesis Kit (NEB #E0554), with primer sequences provided in Table 1. All mutations were confirmed by Sanger sequencing. The hSRF plasmid was obtained from Origene (RG237323). The IE1 (HCMV) expression plasmid was constructed as previously described^94^. For the generation of the HSV-2 ICP0 expression plasmid a doxycycline inducible lentiviral expression vector was synthetically designed and assembled *de novo* to enable tight, tunable expression of the target gene. The construct was based on a tetracycline responsive element (TRE) promoter driving expression of the transgene under the control of the rtTA3 transactivator system, allowing for doxycycline dependent induction of gene expression. ICP0 from HSV-2 strain 333 was codon optimized for human expression. A Kozak sequence (GCCACC) and V5 epitope tag were added to the N-terminus upstream of the start codon, the entire V5-ICP0 cassette was generated via gene synthesis (Twist Biosciences, San Francisco, CA). This fragment was cloned into the lentiviral expression vector using Seamless DNA Assembly Plus Kit (HY-K1041 MedChemExpress LLC., USA) according to manufacturer ‘s protocol. Colonies were screened by restriction digestion using NheI and MluI to confirm correct insert size and orientation. Positive clones were further validated by full-length plasmid sequencing using Sanger sequencing (Microsynth AG, Switzerland). Cells transfected with the HSV-2 ICP0 plasmid were directly co-treated with 100 ng/ml doxycycline to induce expression. The VZV ORF61 expression plasmid^95^ was a kind gift of Jan Rehwinkel (University of Oxford). The KSHV ORF75 expression plasmid was generated as previously described^96^. The cDNA of EBV BNRF1 was amplified using genomic virus DNA as a template and inserted in pcDNA3.1 in frame with a C-terminal myc epitope by Gibson Assembly.

**Table 1:**
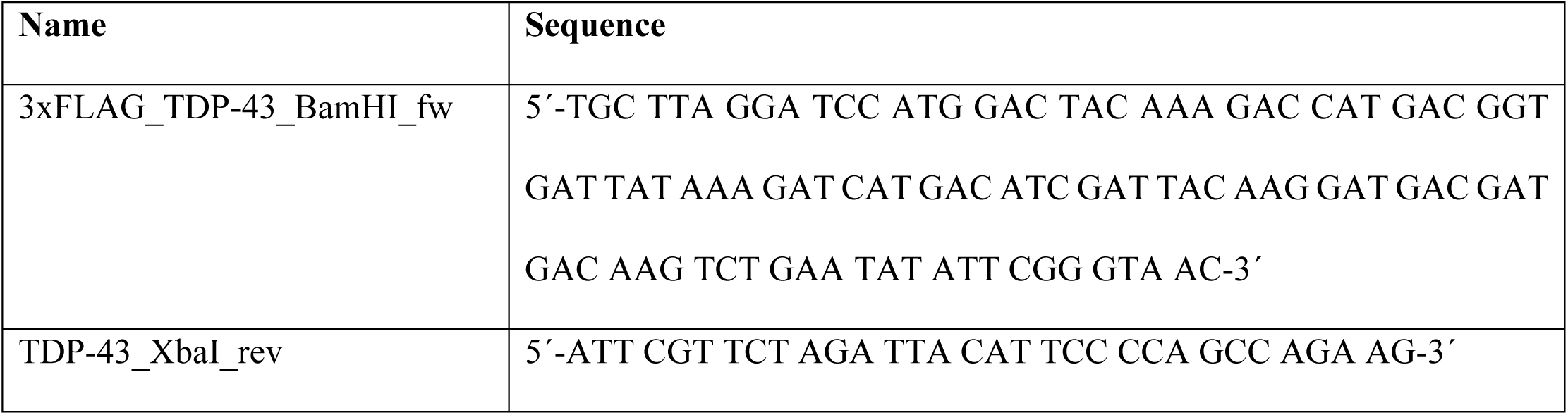
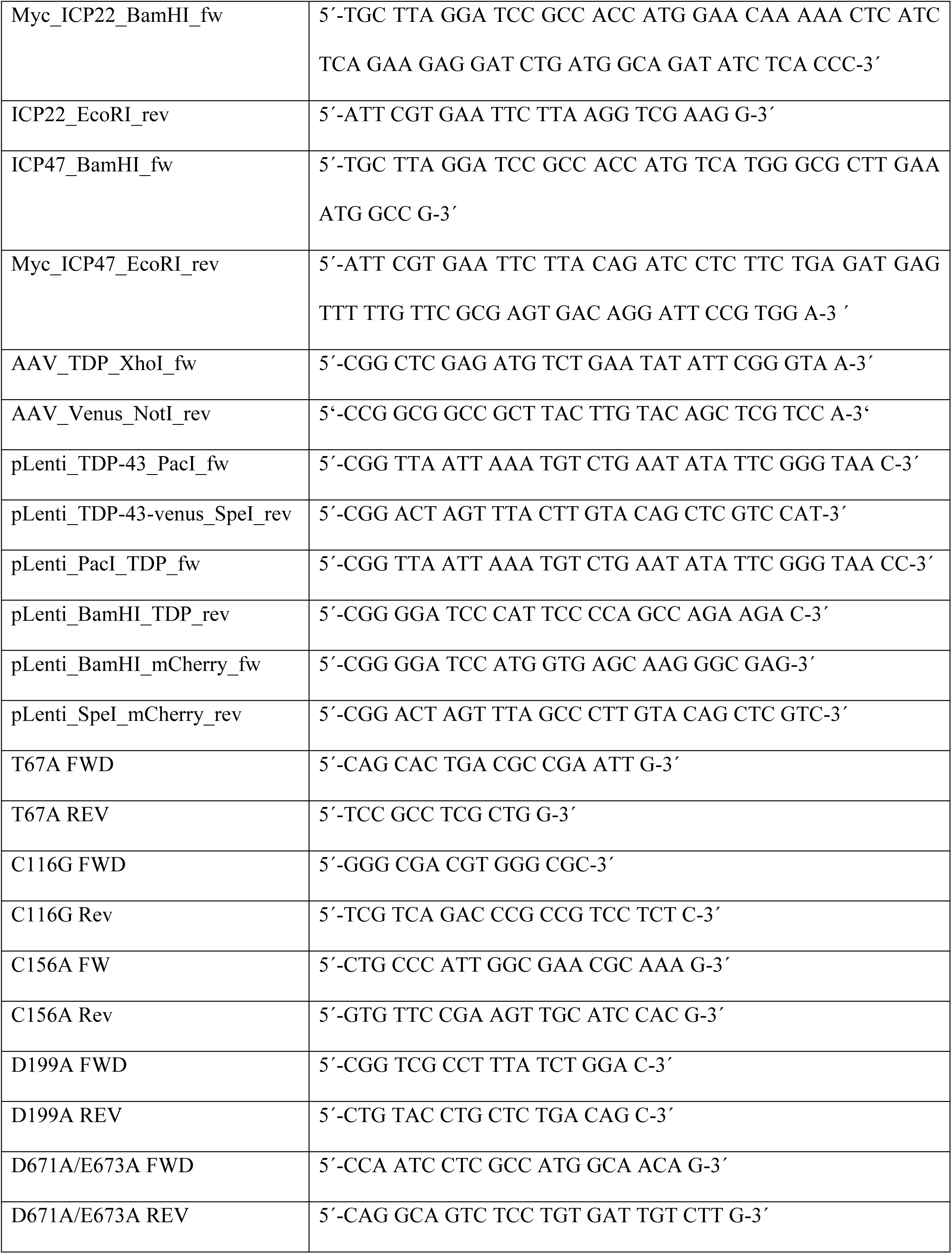
Primer and respective sequence.

#### Infection of forebrain organoids

On day 100, mature forebrain organoids were infected with 20 µl of HSV-1 GFP (3.15×10^5^ FFU/ml) overnight. After virus inoculation, the organoids were washed twice with PBS and fresh neural maturation medium was added. 48 hours post-infection, the media was removed and the organoids were washed twice in PBS and fixed overnight in 4% paraformaldehyde (PFA) at 4 °C.

#### Cryosectioning and immunostaining of forebrain organoids

The day after fixation, organoids were washed twice in PBS and incubated in 30% sucrose solution for 24 h at 4 °C. After dehydration, the organoids were embedded in Tissue-Tek O.C.T., rapidly snap-frozen in dry ice/ethanol and stored at –80 °C. For immunohistochemistry, organoids were cut into 20 μm thick sections and mounted onto superfrost microscope slides. A barrier was created around the sections using a hydrophobic barrier pen (ACD PN 310018) and sections were washed three times with PBS. The sections were first blocked with 5% (v/v) BSA (Gibco 10,270,106) + 0.1% (v/v) Triton X-100 (Sigma-Aldrich, T8787) for 1 h at RT, then blocked with Cohn II γ-globulin blocking solution (Sigma-Aldrich, G2388, 2 mg/ml) for 1 h at 37 °C, 5% CO_2_. Next, the slides were incubated with primary antibody (see below) diluted in 5% (v/v) BSA + 0.1% (v/v) Triton X-100 overnight at 4 °C. After washing three times with PBS for 10 min at RT, sections were incubated in secondary antibody (see below) diluted in 5% (v/v) BSA (Gibco 10,270,106) + 0.1% (v/v) Triton X-100 for 2 h at RT. The sections were washed once with PBS and stained with 4’,6-diamidino-2-phenylindole (DAPI, 1 mg/l; Invitrogen, D1306) diluted in PBS for 10 min at RT. The slides were washed two times with PBS. Sections were mounted using Mowiol mounting medium (Mowiol 4–88 10% (w/v, Carl Roth, 0713), glycerol 25% (w/v, Sigma-Aldrich, G5516), H2O 25% (v/v), 0.2 M Tris HCl pH 8.5 50% (v/v, AppliChem GmbH, A2264), DABCO 2.5% (w/v, Carl Roth, 0718)).

#### Primary Antibodies

Polyclonal rabbit anti-TDP-43 antibody (proteintech, 10782-2-AP, 1:500). Secondary Antibodies: DAPI (Sigma-Aldrich, D9542, 1:1000), goat-rabbit AF-546 (Invitrogen, A11010, 1:500).

#### Immunofluorescence analysis (3D cell culture)

Forebrain organoid images were acquired as 10-15 µm Z-stacks with an interval of 0.5 µm using a Zeiss LSM980 confocal microscope. Analysis of TDP-43 puncta was performed using Imaris (Oxford Instruments, version 11.0.0). For analysis of TDP-43 puncta per cell per image, a surface was created based on the DAPI staining. Based on the DAPI surface, the TDP-43 signal was masked, setting the voxel intensity outside the nucleus to zero. The resulting channel was used for spots creation. The workflow was applied as batch to all images of the same experiments, providing the numbers of nuclei and TDP-43 puncta per image. The number of TDP-43 spots was divided by the number of DAPI surfaces. Thresholds were determined for each experiment individually. To evaluate the number of TDP-43 puncta in GFP positive and negative cells, a surface was created based on the GFP signal. The TDP-43 and DAPI channels were masked based on the GFP surface, setting the voxel intensity outside the surface to zero to create a TDP-43 channel for the infected cells. Additionally, the voxel intensity of the TDP-43 and the DAPI channel was set to zero inside the GFP surface, providing the TDP-43 and DAPI signal for GFP negative cells. Based on the resulting TDP-43 channels, spots were created. Based on the resulting DAPI channels, surfaces were created. The workflow was applied as batch to all images of the same experiments and the number of TDP-43 spots was divided by the number of DAPI surfaces in GFP positive and negative cells. Thresholds were determined for each experiment individually.

#### Immunofluorescence (2D Cell culture)

H4 cells were grown on coverslips (VWR) and treated as indicated. Cells were fixed using 4% (w/v) PFA (SantaCruz) for 10 min. Cells were seeded on PDL-coated glass coverslips in 24 well plates. Following the respective experimental procedure, the cells were washed twice with ice-cold PBS (Gibco, 14190144) and fixed in 4% paraformaldehyde (PFA; Santa Cruz Biotechnology, sc-281692) in PBS for 10 min at RT. Next, the cells were washed twice with PBS, permeabilized with 300 µl 0.1% (v/v) Triton X-100 (Sigma-Aldrich, T8787) + 100 mM Glycine (Sigma-Aldrich, 33226) for 10 min, and blocked with 1.5% (v/v) BSA (Gibco 10,270,106) + 0.1% (v/v) Tween20 (Sigma-Aldrich, P9416) for 1 h at RT. Additionally, 300µl of Cohn II γ-globulin blocking solution (Sigma-Aldrich, G2388, 2 mg/ml) was added and incubated for 1h at 37 °C, 5% CO2. After washing two times with 500 µl PBS, the primary antibody (see below) diluted in 200 µl PBS was added and incubated at room temperature (RT) for 1 h. Next, cells were washed two times with PBS. The secondary antibody (see below) and 4’,6-diamidino-2-phenylindole (DAPI, 1 mg/l; Invitrogen, D1306) was diluted in 200µl PBS and incubated at RT for 1 h. The cells were washed two times with PBS. Glass coverslips were mounted on microscopy slides using self-hardening Mowiol mounting medium (Mowiol 4–88 10% (w/v, Carl Roth, 0713), glycerol 25% (w/v, Sigma-Aldrich, G5516), H2O 25% (v/v), 0.2 M Tris HCl pH 8.5 50% (v/v, AppliChem GmbH, A2264), DABCO 2.5% (w/v, Carl Roth, 0718)). For p-TDP-43 staining the FlexAble 2.0 antibody labelling kit (proteintech) was used according to the manufacturer protocol. After incubation at RT overnight, the Mowiol hardened and over 20 randomly selected images were acquired per sample using a Zeiss LSM980 confocal microscope. Primary Antibodies: Polyclonal rabbit anti-TDP-43 antibody (proteintech, 10782-2-AP, 1:500), mouse anti-lamin B (proteintech, 66095-1, 1:1000), anti-PML (FORTIS Life Sciences, A301-167A, 1:1000), anti-PML (FORTIX Life Sciences, A301-168A, 1:1000), mouse anti-ICP0 (abcam, AB6513, 1:300), mouse anti-ICP27 (Santa Cruz, sc-69806, 1:500), mouse anti-ICP4 (abcam, AB6514, 1:400), rabbit anti-ICP47 (CusaBio, CSB-PA360979ZA01HWY, 1:500), rabbit anti-phospho-TDP-43 (proteintech, 22309-1-AP, 1:500), monoclonal mouse anti-FLAG-tag antibody, (M2, Sigma-Aldrich, F1804, 1:500), mouse monoclonal anti-Myc-tag antibody (Cell Signaling, 2276S, 1:500), mouse monoclonal anti-V5-tag antibody (Cell Signaling, 80076S, 1:500), NeuroTrace 647 (Invitrogen, N21483, 1:100), mouse anti-MAP2 antibody (Invitrogen, 13-1500, 1:400), rabbit IgG control (proteintech, 30000-0-AP, in proportion to the amount of the respective rabbit antibody). Secondary Antibodies: DAPI (Sigma-Aldrich, D9542, 1:10,000), goat anti-rabbit AF-647 (Invitrogen, A21244, 1:500), goat anti-rabbit AF-546 (Invitrogen, A11010, 1:500), goat anti-mouse AF-568 (Invitrogen, A11004, 1:500), goat anti-mouse AF-647 (Invitrogen, A21235, 1:500), goat anti-mouse AF-546 (Invitrogen, A11003, 1:500), FlexAble 2.0 CoraLite Plus 555 Antibody Labeling Kit for Mouse IgG1 (Proteintech, KFA522), FlexAble 2.0 CoraLite Plus 647 Antibody Labelig Kit for Rabbit IgG (Proteintech, KFA503)

#### Immunofluorescence analysis (2D Cell culture)

Foci analysis of immunofluorescence images was performed with Foci Analyzer (v1.91) (publicly available: https://github.com/BioImaging-NKI/Foci-analyzer) in FIJI (ImageJ 1.54p)^97^. In brief, nuclei were automatically defined using DAPI staining and following settings: “StarDist nuclei segmentation 2D” with exclusion of nuclei with a diameter smaller than 4 and larger than 50 µm as well as nuclei located on image edges. The “Foci detection settings” were kept consistent within one experiment but was individually adjusted for each replicate. Following measurements were excluded from the analysis: nuclei with “cell area 2D (µm²)” below 50, nuclei with a mean cell intensity in TDP-43venus (ch1) <5,000, and sum cell intensity (ch1) >10^8^. Further a foci count ch1 = 0 was excluded. To analyse the primary murine neurons and iPSC derived motoneurons, a manually adapted threshold was set in FIJI (ImageJ 1.54p) for each biological repeat, and kept consistent in the analysis between images and infected/uninfected. Afterwards, a region of interest (ROI) was drawn around the cell and particles were counted within the ROI.

#### Live cell imaging localization analysis of TDP-43venus

H4 cells were seeded in 96-well plate and one day after transduced with AAV expressing TDP-43venus for 3 days. Afterwards, HSV-1 infection at an MOI 0.3 for 17 h. Images were acquired using the Incucyte SX5 (Sartorius). Images were analysed using Imaris (Oxford Instruments, version 11.0.0). A surface was created based on the DAPI signal. TDP-43venus signal in the nucleus was masked. Using the resulting channel, a surface of the extranuclear TDP-43venus was created. The sum intensity of the extranuclear TDP-43venus fluorescence was divided by the number of DAPI surfaces per image.

#### Transient transfection, LT1

H4 cells were seeded to reach a confluency of 60–80% for transfection. Plasmid DNA was suspended in Opti-MEM (Gibco 31985070), and after 5 min TransIT-LT1 (Mirus, MIR 2300) (3 µl per 1 µg plasmid DNA) was added. After incubation at RT for 20 min, the transfection mix was added to each well. Medium was replaced with fresh medium after 6–16 h.

#### Transient transfection, calcium phosphate

HEK293T or Lenti-X 293T cells were transiently transfected at 80% confluency one day after seeding, using respective plasmids. Plasmid DNA was adjusted according to the culture format: for 12-well plates, 2 µg DNA was diluted in 25 µl ddH₂O, mixed with 4 µl 2.5 M calcium chloride and 40 µl 2× HBS; for 10 cm dishes, 15 µg DNA was diluted in 450 µl ddH₂O, mixed with 25 µl 2.5 M calcium chloride and 250 µl 2× HBS. After incubation for 3 min at RT, 2 ml (12-well plate) or 9 ml (10 cm dish) DMEM(++) was added. The culture medium of the seeded cells was aspirated and replaced with the transfection mixture. 16 h post-transfection, the medium was exchanged for fresh DMEM(++).

#### Solubility assay

H4 cells (0.8 x 10^6^ cells per dish) were seeded in 10 cm dishes, transduced with AAV TDP-43venus (MOI 1000), and infected with HSV-1 (MOI 0.3). 17 h after infection, cells were washed with 2.5 ml dPBS and harvested in 200 µl RIPA buffer. Protein concentrations were determined by Bradford assay and adjusted to 1 mg/ml, with 20 µl taken as whole cell lysate (WCL). Samples (100 µl) were centrifuged at 100,000 × g for 30 min at 4°C. The supernatant (SN, 20 µl taken as soluble fraction S) was separated, and the pellet was resuspended and sonicated in 200 µl RIPA buffer, followed by another 100,000 × g spin for 30 min at 4 °C. The final pellet was dissolved and sonicated in 100 µl 8 M Urea buffer, with 20 µl collected as insoluble fraction (I). Samples WCL, S, and I were mixed with 10 µl 4X sample buffer and boiled at 95 °C for 5 min. Samples were subjected to SDS-PAGE and Western blot analysis.

#### Nuclear and cytoplasmic fractionation

H4 cells (0.8 x 10^6^ cells per dish) were seeded in 10 cm dishes, transduced with AAV TDP-43venus (MOI 1000) for 3 days, and then partially infected with HSV-1 (MOI 0.3). 17 h after infection, cell culture medium (DMEM(++)) was removed and cells washed two times with ice-cold PBS. Then, cells were removed with a cell scraper, collected in 1 ml of ice-cold PBS and stored in a microcentrifuge tube. The tube was centrifuged for 10 s. with a table top centrifuge and the supernatant discarded. The cell pellet was dissolved in 1 ml of ice-cold 0.1% NP40-PBS (NP40, Calbiochem). An aliquot of 200 µl was taken and stored in a separate tube as the whole cell lysate (WCL). The rest of the sample was then centrifuged for 10 s. with a table top centrifuge. Supernatant was removed and stored as the cytoplasmic fraction (C). The remaining cell pellet was resuspended in 1 ml of ice-cold 0.1% NP40-PBS and again centrifuged for 10 s. with a table top centrifuge. Supernatant was discarded and the white pellet stored as the nuclear fraction (N). Samples were then applied to Western blot for further analysis.

#### SDS-PAGE and western blot analysis

Samples were applied to SDS polyacrylamide gel (12% separation gel, 5% loading gel) inserted in SDS-PAGE chamber containing SDS-running buffer (25 mM Tris, 192 mM Glycine, 1 g/L SDS). Electrophoresis was performed in two steps: first at 80 V for 20 min and then at 100 V for ∼3 h. Using the western blotting system by Invitrogen Corporation (XCell II Blot Modul EI0002) containing blotting buffer (25 mM Tris, 192 mM Glycine, 20% methanol) samples were transferred to a Nitrocellulose membrane (0.2 µm, Amersham Protran, 10600011) for 1.5 h at 25 V. The membranes were then blocked in 5% milk powder in 1x TBS + 0,05% Tween20 (Sigma-Aldrich, P9416) for 1 h at RT. Afterwards, the membranes were incubated with primary antibodies (see below), diluted in 1x TBS-T overnight at 4 °C. The membranes were washed 3 times with TBS-T, before incubation with secondary antibodies (see below), diluted in 5% milk at RT for 2 h. Protein bands were detected by adding Blotting Substrate (Thermo Fisher Scientific, 34577) to the membrane and imaged via the Odyssey M (Li-cor). Western Blot analysis of immunoprecipitation assay samples of SUMOylated proteins were blotted as described above onto Immun-Blot PVDF membrane (0.2 µm, Bio-Rad, 1620177). Membranes were blocked in 5% Bovine serum albumin (BSA, Carl Roth, 8076.2) in TBS-T. Antibodies for SUMOylation immunoblots were incubated in 1% BSA in TBS-T.

#### Primary antibodies

Monoclonal mouse anti-FLAG-tag antibody, M2 (1:5,000), Sigma-Aldrich, F1804), anti-GAPDH (BioLegend, 607902; 1:1,000), Polyclonal rabbit anti-TDP-43 antibody (proteintech, 10782-2-AP, 1:2,000), mouse anti-lamin B (proteintech, 66095-1, 1:1,000), mouse anti-HSV-1 ICP0 (abcam, AB6513, 1:7,500), anti-SUMO2 (novus biologicals, NBP1-77163, 1:1,000) Secondary Antibodies: anti-rabbit IgG HRP (Promega, W401B, 1:10,000), anti-mouse IgG HRP (Promega, W402B, 1:10,000), Veriblot HRP (abcam, ab131366, 1:1,000)

#### Luciferase assay

HEK293 cells were seeded in 12-well plates (1×10⁵ cells/well) and transfected with pcDNA3, TDP-43-L1 and TDP-43-L2 using calcium-phosphate precipitation. The medium was replaced with Opti-MEM the next day, and cells were infected with HSV-1 (MOI 0.3) or mock treated. After 48 h post-infection cells were lysed in passive lysis buffer (Promega, E1941). 50 µl of each sample was transferred into a white Nunc F-bottom 96-well plate. To measure Gaussia luciferase activity the plate was put into an Orion II Microplate Reader (Berthold). Automatically 50 µl of a 20 µM coelenterazine (PJK biotech) stock solution was added and after 2 s the relative light units per second (RLU/s) were measured for a period of 1 s.

#### Fluorescence recovery after photobleaching

H4 cells were seeded at a density of 10,000 cells per 35 mm ibidi dishes. After 24 h, cells were transduced with lentiviruses expressing TDP-43halo. After 3 days, cells were infected with HSV-1 GFP with an MOI of 0.3. After 17 hours, cells were washed 3 times with PBS and the TDP-43halo was stained using the Janelia Fluor 646 HaloTag Ligand (Promega, HT1060) according to the manufacturer’s instruction. Afterwards, the medium was changed to DMEM without phenol red and the dishes were placed in a stage-top live-cell incubation chamber (37 °C, 5% CO_2_) of a Leica DMi8 confocal microscope. Images were acquired using the HC PL APO CS2 20x/0.75 IMM objective. Pre-/Post-bleaching laser settings were kept consistent. Pre-bleaching two images were taken (2 s interval). During bleaching, the intensity of laser line (633 nm) was increased to 100% for 5 s and solely applied to the region of interest (ROI). Post-bleaching, images were taken every 5 s for 50 s. The ROI was confined to a single dot like structure in the infected samples. For the mock control an ROI of similar size was taken. The analysis was performed using the Leica Application Suite X 3.7.6.25997. The mean fluorescence intensity (MFI) of the ROI was quantified at each time point and the MFI pre-bleaching was set to 100% and post-bleaching to 0%.

#### Holographic and fluorescent imaging (NanoLive)

H4 cells (35,000 cells/dish) were seeded in a 35 mm µ-Dish (ibidi, 80136) closed with a DIC Lid for µ-Dishes (ibidi, 80051). The next day, cells were transduced with TDP-43mCherry expressing lentiviruses. After 3 days cells were washed twice with PBS and medium was exchanged to DMEM without phenol red. Afterwards, cells were infected with HSV-1 GFP (Strain F) with an MOI 0.3. Subsequently, dishes were placed into the top-stage incubator of the 3D cell explorer (NanoLive) at 37 °C, 5% CO_2_. Imaging started 4 h post-infection, acquiring images every 15 min, using the refractive index (RI, holographic reconstruction), FITC and TRITC fluorescence channels (Eve, Version 3.0.2.59).

#### Immunoprecipitation assay of SUMOylated proteins

HEK293T cells were transiently transfected with plasmids encoding for TDP-43-FLAG and/or HSV-1 ICP0 or an empty control for 48 h. Cells were treated with sodium arsenite (1 h, 500 µM). Cells were harvested and lysed using following buffer 150 mM Tris-HCl (pH 6.7), 5% SDS, 30% Glycerol, 20 mM NEM. Briefly, the cells were harvested in PBS supplemented with 20 mM N-ethylmaleimide (NEM, MedChemExpress, HY-D0843). Cells were lysed in SDS lysis buffer supplemented with 20 mM NEM and protease inhibitor cocktail (PI, Sigma-Aldrich, 11697498001). Lysates were 10-fold diluted in PBS supplemented with 0.5% NP-40 (Biozol, USB-N3500), 20 mM NEM and PI (1:500), pulse-sonicated 3 times for 5 s each and the supernatants cleared for 10 min at 12,000 g at 4 °C. 10% of the supernatants was treated with benzonase (5 U, Biozol, PTG-PX-P4894-25000U) and kept as input samples. The residual supernatant was incubated with pre-washed and pre-coated anti-FLAG magnetic beads (Merck, M8823) for 4 h at 4 °C on a rotary shaker. The beads were washed 3 times with washing buffer (10 mM Tris-HCl, pH 7.5, 150 mM NaCl, 0.05% NP40, 0.05 mM EDTA, 20 mM NEM, PI 1:500) before resolving the proteins by SDS-PAGE and Western Blotting.

#### MG-132 treatment

H4 cells were seeded (10^4^ cells per well) on PDL-coated glass slides in a 24 well plate, transduced with lentiviruses expressing TDP-43venus (1 µl per well) on the next day and incubated for 3 days in culture (37 °C, 5% CO_2_). Cells were washed with PBS three times; new medium was added (DMEM(++)) and infected with HSV-1 with an MOI of 0.3. The proteasome inhibitor MG-132 (Hycultec, HY-13259) was added 1 h after infection at a concentration of 3 µM. Cells were fixed with 4% PFA 17 h after infection.

#### *In vivo* HSV-1 infection

Mice of both sexes were infected with HSV-1 as previously described at the animal BSL-2 (ABSL-2) facilities of CC-FRIC^98^. Briefly, mice (10-to-16-week old, age-and sex-matched across treatments) were anesthetized and then intracerebrally inoculated with HSV-1 (KOS strain) in sterile PBS at the indicated PFU or with PBS (mock infection). Mice were daily monitored for weight loss and disease progression/clinical scores according to Institutional Animal Care and Use Committee (IACUC) guidelines. Blood samples were collected at the indicated times post-infection via retro-orbital bleeding, and sera were immediately separated and stored at –80 °C. Mice were humanely euthanized at the indicated times post-infection for brain tissue collection or euthanized following humane endpoint criteria as defined by IACUC guidelines.

#### Enzyme-linked immunosorbent assay (ELISA)

NF-L protein levels in the sera of mock-treated or HSV-1-infected mice were quantified by ELISA (Novus Biologicals, Cat # NBP2-80299) following the manufacturer’s instructions.

#### Cryo-sectioning and immunofluorescence of mouse brain

Whole brain tissues were divided into half (mid-sagittal sections) and fixed with 4% (v/v) PFA in PBS for 24 h. Next, brain sections were washed twice in PBS and then incubated in 30% (v/v) sucrose in PBS for 24 – 48 h at 4 °C. After dehydration and sucrose protection, the brains were embedded in Tissue-Tek O.C.T. compound, snap-frozen in dry ice, and stored at –80 °C. Brains were cut into 15 μm thick mid-sagittal sections, mounted onto superfrost microscope slides, and subjected to immunofluorescence staining as described above for human brain organoids. The primary antibodies and dilutions used for the immunostaining of brain sections are rat monoclonal anti-p-S409/410-TDP-43 (Biolegend, 829901, 1:200) and goat polyclonal anti-HSV-1 (Invitrogen, PA1-7493, 1:1000). The following secondary antibodies were used: AlexaFluor 488 donkey anti-goat (Invitrogen, A11055, 1:1000), and AlexaFluor 488 donkey anti-goat (Invitrogen, A48272, 1:1000).

#### H&E and immunohistochemistry

Formalin-fixed paraffin-embedded (FFPE) tissue blocks were sectioned at 4 μm and subjected to hematoxylin–eosin (H&E) staining using the HE 600 automated stainer (Ventana Medical Systems Inc.) according to the manufacturer’s instructions. Immunohistochemical staining was performed on an automated, validated, and accredited staining platform (Ventana Benchmark ULTRA, Ventana Medical Systems Inc.) using the OptiView Universal DAB Detection Kit. After automated deparainization, heat-induced antigen retrieval was performed with CC1 solution (#950-500, Ventana) for 32 min. Sections were then incubated with anti-phospho-TDP-43 (pS409/410) mouse monoclonal antibody (clone 11-9, Cosmo Bio; dilution 1:5.000) for 32 min at 37 °C. Detection was performed using the OptiView system, followed by hematoxylin II counterstaining for 8 min and blue colouring reagent for 8 min, in accordance with the manufacturer’s guidelines. Stained slides were scanned using a NanoZoomer HT 2.0 digital slide scanner (Hamamatsu Photonics, Japan).

#### Infection inhibition assay

H4 cells (10,000 cells /well) were seeded in a 96-well plate and transduced the next day, using AAV expressing TDP-43venus. After 3 days, cells were infected with either HSV-1 or HSV-1 simultaneously co-treated with an infection inhibiting antibody (200 µg/ml) at an MOI of 0.3. The antibody is a chimeric monoclonal antibody hu2c purified from serum-free SP/0 cell supernatants by chromatography as described previously^99,100^. Images were acquired hourly with a 20x objective and eventually analysed using the Incucyte SX5 Live-Cell Analysis System (Sartorius).

#### Pseudoparticle production

To produce pseudotyped VSVΔG-GFP particles, 6 × 10⁶ HEK293T cells were seeded 18 h before transfection in 10 cm dishes. Cells were transfected with 15 µg of the glycoprotein-expression vector using polyethylenimine (PEI; 1 mg/ml in H₂O; Sigma-Aldrich). 24 h post-transfection, cells were infected with VSVΔG-GFP particles pseudotyped with VSV-G at a multiplicity of infection (MOI) of 3. 1 h post-infection, the inoculum was removed. Pseudotyped VSVΔG-GFP particles were harvested 16 h post-infection. Cell debris was pelleted and removed by centrifugation (500 × g, 4 °C, 5 min). Residual input particles carrying VSV-G were neutralized by adding 10% (v/v) I1 hybridoma supernatant (I1; mouse hybridoma supernatant from CRL-2700, ATCC) to the cell culture supernatant.

#### Serum inhibition

HEK293T cells (1 × 10⁵/well) were seeded in 96-well plates and infected with 1×10⁴ VSVΔG-GFP pseudoparticles carrying the corresponding viral glycoprotein (GP), which had been preincubated (30 min, 37 °C) with the indicated volume of serum. 24 h post-infection, GFP-positive cells were automatically quantified using a Cytation 3 microplate reader (BioTek).

#### HSV-1 kinetics

H4 cells were transduced using AAV expressing TDP-43venus. Three days later, cells were infected with HSV-1 with an MOI of 0.3. Infection was monitored using a Cytation 3 microplate reader (BioTek), and particle counts as well as size were determined using ImageJ.

#### Acyclovir treatment

Cells were infected with HSV-1 with an MOI of 0.3, and treated with Acyclovir (ACV, Ratiopharm, 1 mM) after 4 and 7 h post-infection. Cells were then incubated for another 3 days before fixation with 4% PFA for 10 min.

#### Software

For statistical analysis of the electronic health records and serology data, R (v.4.5.2) was used with packages ‘stats’ (v4.5.2) and ‘tidyverse’ (v2.0.0). For visualization of quantification and statistical analyses GraphPad PRISM 10 was used. P-values were calculated using t test, Wilcoxon matched-pairs signed rank test, Kruskal-Wallis test followed by Dunn’s multiple comparisons test, Spearman’s rank correlation test. Unless specified otherwise, data are shown as the mean of at least three biological replicates ± SEM. Significant differences are indicated as: *p < 0.05; **p < 0.01; ***p < 0.001. Insignificant differences are not indicated.

## DATA AVAILABILITY

All data is available upon reasonable request to the corresponding authors. Source data are provided with this paper.

## ACKNOWLEDGEMENTS

We thank Ramona Bück, Jana-Romana Fischer, Birgit Ott, and Pia Veratti for amazing technical assistance. Recombinant AAV was provided by the Core Facility Viruses of the Medical Faculty Ulm. We acknowledge funding by the German Federal Ministry of Education and Research (BMFTR; IMMUNOMOD-01KI2014 to K.M.J.S. and Duxdrugs-01K17 to F.F.), the German Research Foundation (DFG; CRC1279, CRC1506, SP 1600/13-1, SP 1600/7-1, SP 1600/9-1). K.M.J.S. and K.M.D. acknowledge funding by the DZNE Stiftung. K.M.D was funded by the German Research Foundation (DFG; CRC1506 and CRC1149), and received support from Target ALS Grant No. BM-2024-C2-L4. T.S. was funded in the framework of the Research Unit FOR5200 DEEP_DV (443644894) through project STA357/8-2. This work was further supported by the ‘Where There is Light Foundation’ and institutional funds by Cleveland Clinic Foundation (to M.U.G.). D.F., D.R., Z.E., A.d.L., A.G. and A.P. are part of the International Graduate School for Molecular Medicine, Ulm (IGradU). We dedicate this work to Karl-Klaus Conzelmann (1955-2025) who was an inspirational neurovirologist and mentor.

## ETHICS DECLERATION

The authors declare no competing interests.

## Notes

### Competing Interest Statement

The authors have declared no competing interest.

