## Extended data Figures for "Herpes simplex virus infection promotes ALS pathology through ICP0-mediated PML body disruption"

**Extended Data Fig. 1-5**

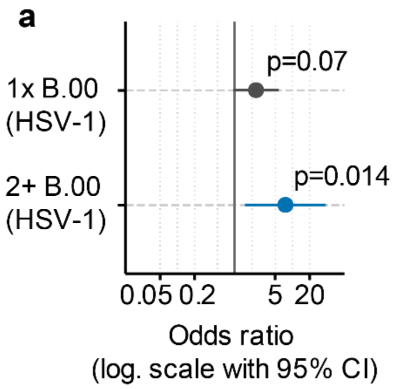


**b**

| Characteristic | N = 1,134[1] |
|---|---|
| Data | |
| Complete | 937 (83%) |
| Incompl. data | 197 (17%) |
| CMV_Status | |
| neg | 467 (43%) |
| pos | 611 (57%) |
| equivocal | 56 |
| HSV1_Status | |
| neg | 171 (15%) |
| pos | 953 (85%) |
| equivocal | 10 |
| VZV_Status | |
| neg | 124 (13%) |
| pos | 839 (87%) |
| equivocal | 171 |
| EBV_Status | |
| neg | 41 (3.6%) |
| pos | 1,089 (96%) |
| equivocal | 4 |

[1] n (%)

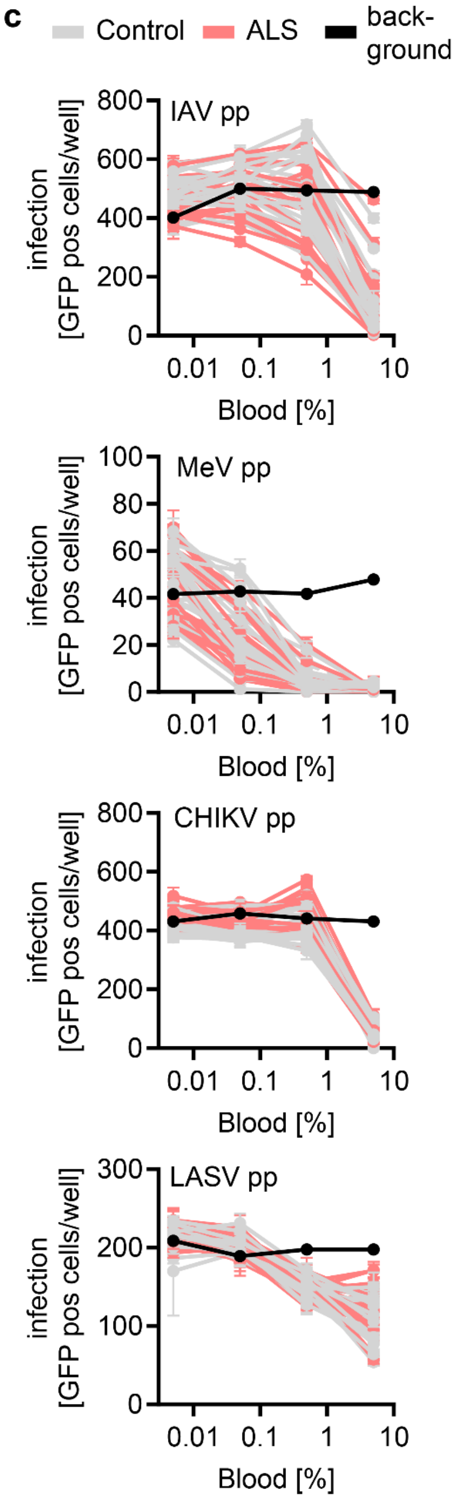


**Extended Data Fig. 1 Serology against other viruses. a**, Conditional logistic regression model demonstrates increased odds for ALS diagnosis with >1 prior HSV-1 calendar quarter diagnoses. **b**, Serology for CMV, HSV-1, VZV and EBV in the cross-sectional serology cohort (n = 1,134; n = 937/17% samples with complete data for all 4 tests). 57% of the participants were seropositive for CMV, 85 % for HSV-1, 87 % for VZV, and 96 % for EBV. Samples with equivocal results interpretation were excluded. **c**, Inhibition of VSVΔG(GFP) pseudotyped with indicated viral glycoproteins by varying concentrations of sera (x-axis) from control (grey) and ALS patients after symptom onset (red). Black line indicates pseudoparticles without serum. Each line represents one donor and the mean of three biological replicates.

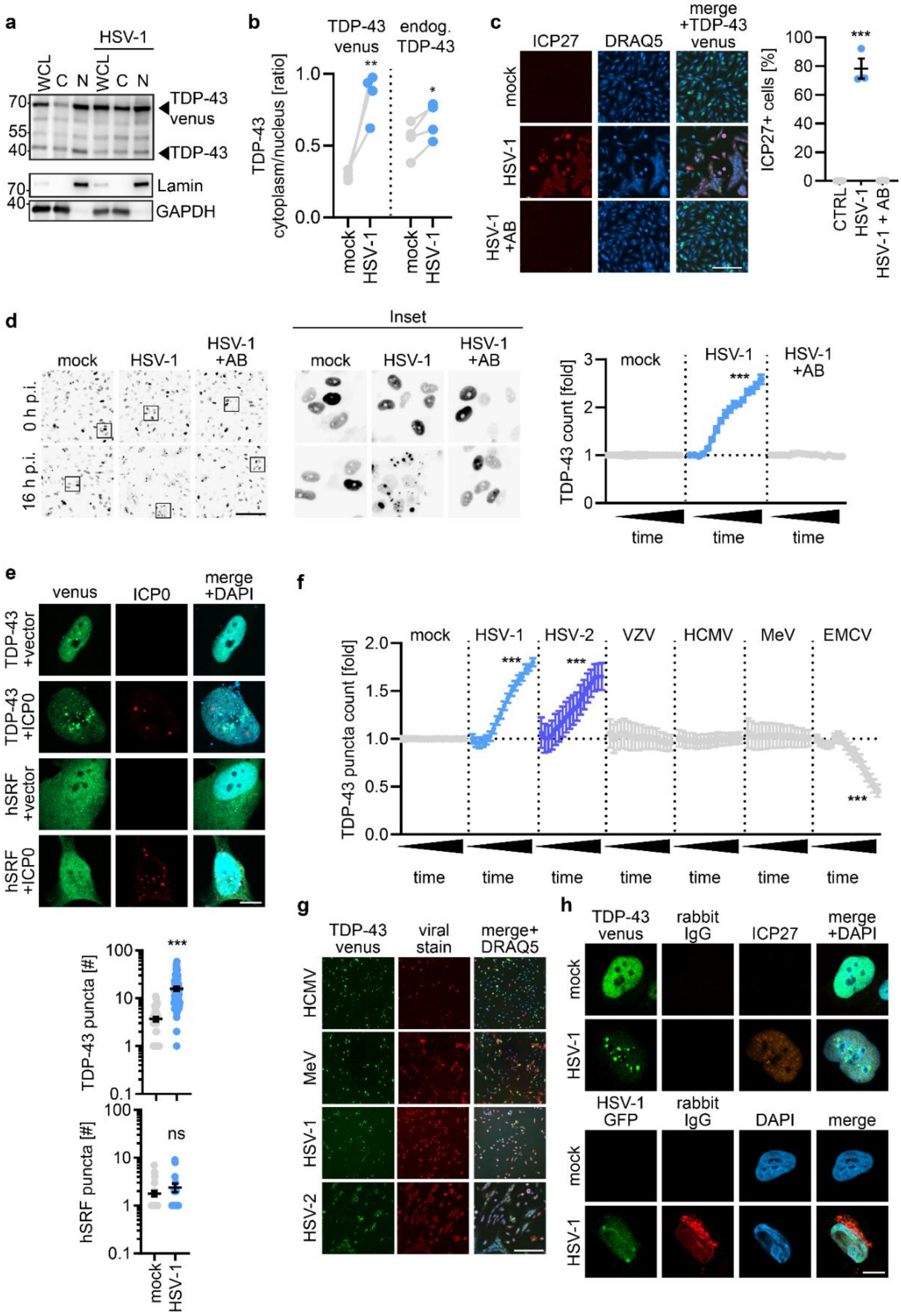


**Extended Data Fig. 2 Specificity of HSV-1 infection towards TDP-43. a**, Immunoblotting of TDP-43 and GAPDH in H4 cells, transduced with AAV-TDP-43venus and after three days infected with HSV-1 for 17 h. Cells were harvested and separated into nuclear (N) and cytosolic (C) fractions. **b**, Ratio of cytosolic to nuclear TDP-43 in (**a**), either overexpressed (TDP-43venus, left) or endogenous (endog., right). n = 4, *p<0.05 **p<0.01, paired t test. **c**, Left, representative immunofluorescence images of AAV TDP-43venus (green) transduced H4 cells

17 h post-HSV-1 (MOI 0.3) infection (ICP27, red) treated with a neutralizing antibody (200 µg/ml). DRAQ5 (blue) Scale bar, 200 µm. Right; quantification of ICP27 positive cells divided by nuclei count (DRAQ5). Lines represent mean±SEM, n = 3, three independent biological replicates, ***$p<0.001$, t test. **d**, Left, representative start and endpoint images of TDP-43venus kinetic of AAV TDP-43venus transduced H4 cells infected with HSV-1 (MOI 0.3) and treated as in (**c**). Images were acquired every hour over a time period of 16 h. Black squares indicate origin of inset. Scale bar, 200 µm. Right, quantification of the kinetic. The number of TDP-43venus particles was normalized to the start of the measurement and fold of the control (mock) at respective time point. One biological replicate. A linear regression model with time and treatment as fixed effects demonstrates a significant effect for treatment (***$p<0.001$) and treatment:timepoint interaction (***$p<0.001$). Pairwise comparison with estimated marginal trend contrasts compared to the control group or to HSV-1+AB demonstrates a significant difference of HSV-1 (***$p<0.001$) **e**, (upper panel) H4 cells co-transfected with either TDP-43venus (green) and ICP0 (red) or human serum response factor-GFP (hSRF-GFP, green) and ICP0 (red). Cells were fixed after 24 h and imaged. DAPI (blue). (lower panel) Quantification of TDP-43venus and hSRF-GFP puncta. Lines represent mean±SEM, n = 25-58, three independent biological replicates, ***$p<0.001$, t test. **f**, H4 cells transduced with AAV TDP-43venus and three days later infected with indicated virus with an MOI of 0.3. Images of TDP-43venus were acquired hourly for 17 h and analyzed regarding puncta formation. Each count was first normalized to the count of the start of the measurement (0 h post-infection) and then to respective mock count fold. Three biological replicates. A linear regression model with time and treatment as fixed effects demonstrates a significant effect for treatment (***$p<0.001$) and treatment:timepoint interaction (***$p<0.001$). Pairwise comparison with estimated marginal trend contrasts compared to the control group demonstrates a significant difference of HSV-1 and HSV-2 (***$p<0.001$) **g**, Endpoint (17 h post-infection) immunofluorescence staining of viral proteins (HCMV IE1, MeV-RFP RFP, HSV-1 ICP27, HSV-2 ICP27) in red to confirm successful infection in (f). DRAQ5 (blue). Scale bar, 200 µm. **h**, Isotype control (IgG rabbit derived) staining of HSV-1 infected H4 cells (MOI 0.3, 17 h). Upper panel; transduced with AAV TDP-43venus (green) and infected with HSV-1 H4 cells were blocked using standard blocking protocol (1.5% (v/v) BSA, 1 h, RT) and additionally with Cohn II γ-globulin blocking solution. IgG-rabbit (red), ICP27 (orange), DAPI (blue). Lower panel; H4 cells infected with HSV-1 GFP (green), and blocked using only the standard blocking protocol without Cohn II γ-globulin blocking solution. IgG-rabbit (red), DAPI (blue). Scale bar, 10 µm.

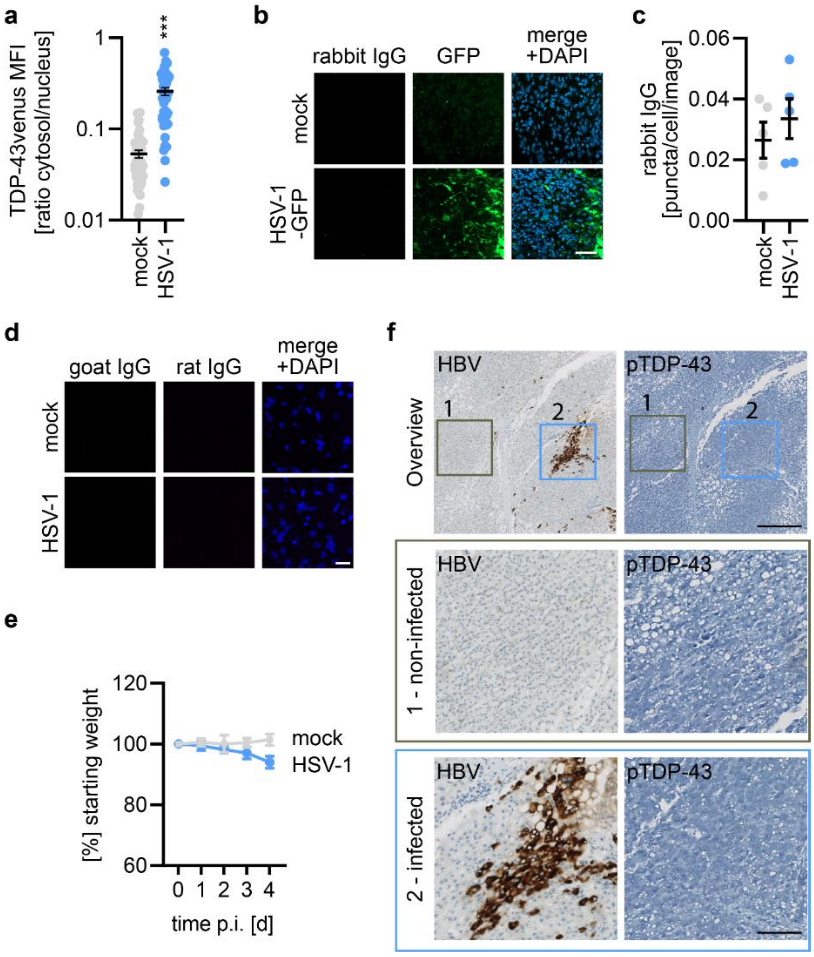


**Extended Data Fig. 3 Impact of HSV-1 infection in human tissue. a**, Ratio of quantified cytosolic and nuclear TDP-43venus mean fluorescence intensity (MFI) in Fig. 3a. Lines represent mean±SEM, n = 41-43, three independent biological replicates. ***p<0.001, t test. **b**, Isotype staining control (IgG rabbit) (purple) of infected (HSV-1 GFP, green) and uninfected organoids displayed in Fig. 3a. **c**, Quantification of isotype control (IC) puncta in (b). **d**, Isotype staining control of HSV-1 (IgG goat, green) and pTDP-43 (IgG rat, purple) in infected and uninfected conditions as in Fig. 3g. Scale bar, 20 µm. **e**, Bodyweight surveillance of mice (Fig. 3k) post-HSV-1 infection (blue) or mock infection (grey). Values are given as percentage of bodyweight on the infection date. Dots represent mean±SEM, n = 3. **f**, Immunohistochemistry staining of a liver tissue sample from a patient with Hepatitis B virus (HBV) infection. Viral marker (hepatitis B surface antigen) in brown and pTDP-43 in purple. Scale bar, 400 µm, scale bar inset, 50 µm.

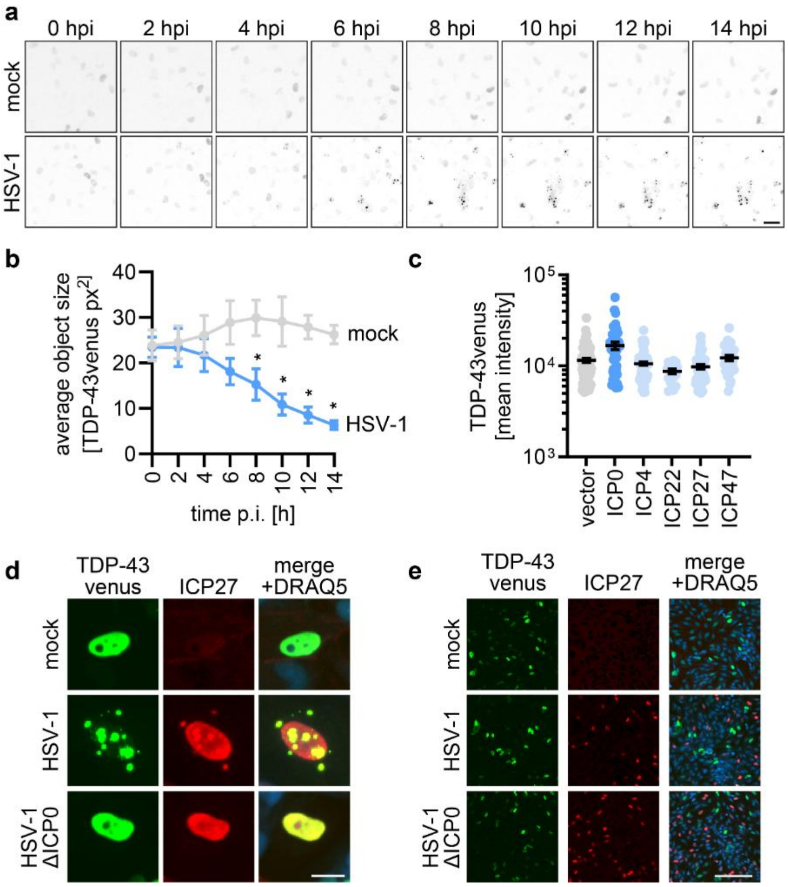


**Extended Data Fig. 4 Influence of HSV-1 on TDP-43venus. a**, Image series of TDP-43venus (AAV) transduced H4 cells either infected (HSV-1 MOI 0.3) or mock treated. Scale bar, 200 µm. **b**, Analysis of the average TDP-43venus object size in (**a**). Dots represent mean±SEM, n = 3, three independent biological replicates, *$p<0.05$, t-test. **c**, Quantification of the mean TDP-43venus intensity of each analyzed nucleus in Fig. 4g. Lines represent mean±SEM, dots represent individual analyzed cells, n = 17-94, three independent biological replicates. **d**, Representative images of H4 cells transduced with AAV TDP-43venus (green) and infected with mock, HSV-1, or HSV-1ΔICP0 for 17 h as in Fig. 4j. ICP27 (red), DRAQ5 (blue). Scale bar, 10 µm. **e**, Immunofluorescence images of H4 transduced with AAV overexpressing TDP-43venus (green). After 3 days, cells were infected with HSV-1 or HSV-1ΔICP0 (MOI 0.3) and fixed 7 h post-infection. Infected cells were assessed via ICP27 (red) staining. DRAQ5 (blue) Scale bar, 200 µm.

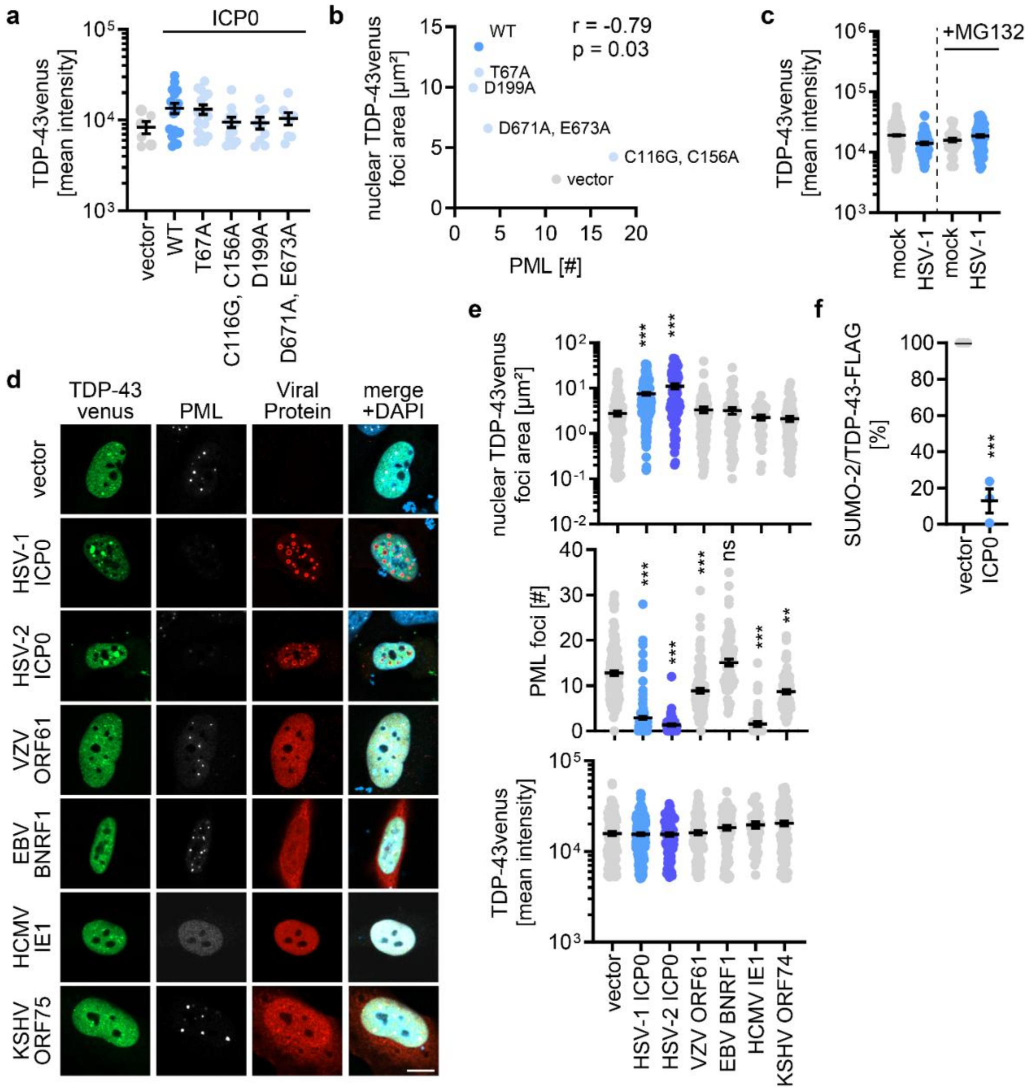


**Extended Data Fig. 5 Impact of HSV-1 ICP0, HSV-1 ICP0 mutants, and viral homologs of ICP0 on TDP-43venus.** **a**, Quantification of the TDP-43venus mean intensity of nuclei analyzed in Fig. 5c. Lines represent mean±SEM, dots represent individual cells, n = 37-74, three independent biological replicates. **b**, Plotting of total nuclear TDP-43venus foci area (y-axis) and respective number of PML foci (x-axis) in Fig. 5c. ICP0 wild-type (WT, blue), ICP0 mutants (light blue) and vector (grey). Dots represent the mean of n = 37-74, three independent biological replicates. **c**, Nuclear TDP-43venus mean intensity of analyzed cells in Fig. 5f. Lines represent mean±SEM, dots represent individual cells, n = 58-362, three independent biological replicates. **d**, Confocal images of co-transfected H4 cells with TDP-43venus (green) and one viral protein (HSV-1 ICP0, HSV-2 ICP0 (+100 ng/ml doxycycline), VZV ORF61, EBV BNRF1, HCMV IE1, KSHV ORF75). Cells were fixed after 24 h of transfection and stained for PML (white), viral protein (red), DAPI (blue). Scale bar, 10 µm. **e**, Quantification of nuclear TDP-43venus foci area (top), PML foci count (mid), and TDP-43venus mean intensity (bottom) of transfected H4 cells in (**d**). Lines represent mean±SEM, n=26-198, three independent biological replicates, ****$p<0.0001$ Kruskal-Wallis test followed by Dunn's multiple comparisons test. **f**, Quantification of the mono-SUMO-2 lane in the TDP-43 immunoprecipitation of Fig. 5g normalized to the vector control. n = 3, ***$p<0.001$ paired t test.
